# A local ATR-dependent checkpoint pathway is activated by a site-specific replication fork block in human cells

**DOI:** 10.1101/2023.03.26.534293

**Authors:** Sana Ahmed-Seghir, Manisha Jalan, Helen E. Grimsley, Aman Sharma, Shyam Twayana, Settapong T Kosiyatrakul, Christopher Thompson, Carl L. Schildkraut, Simon N. Powell

## Abstract

When replication forks encounter DNA lesions that cause polymerase stalling a checkpoint pathway is activated. The ATR-dependent intra-S checkpoint pathway mediates detection and processing of sites of replication fork stalling to maintain genomic integrity. Several factors involved in the global checkpoint pathway have been identified, but the response to a single replication fork barrier (RFB) is poorly understood. We utilized the *E.coli*-based Tus-*Ter* system in human MCF7 cells and showed that the Tus protein binding to *TerB* sequences creates an efficient site-specific RFB. The single fork RFB was sufficient to activate a local, but not global, ATR-dependent checkpoint response that leads to phosphorylation and accumulation of DNA damage sensor protein γH2AX, confined locally to within a kilobase of the site of stalling. These data support a model of local management of fork stalling, which allows global replication at sites other than the RFB to continue to progress without delay.

## Introduction

Genomic DNA is constantly exposed to exogenous and endogenous damaging agents, forming DNA lesions that challenge the progression of replication forks (RFs) during the S-phase of the cell cycle. The genome contains natural impediments, which cause replication stalling such as common fragile sites, repeated sequences, non-B structures, sites of collision between transcription and replication machinery, or DNA-protein complexes(Mirkin and Mirkin, 2007). These replication fork barriers (RFB) can impede the progression of the replication machinery and if unresolved, lead to genomic instability, a common hallmark of aneuploidy, neurological and neuromuscular disorders, and cancer.

Replication stress can impede the progression of the replication fork. During replication stress, the intra-S phase checkpoint pathway can be activated at a local and global level which is a critical step to process the DNA lesions and maintaining genome integrity. The S-phase checkpoint involves the uncoupling of replicative polymerases from the replicative helicase and the generation of ssDNA, which rapidly gets coated with RPA. In turn, this activates ATR/ATRIP kinase signaling with its effector kinase CHK1, plus phosphorylation of H2AX at Ser139 in response to a variety of lesions which promote cell cycle arrest, fork stabilization, and restart(Iyer and Rhind, 2017, 2013; Willis and Rhind, 2009a; Zeman and Cimprich, 2013).

Previous studies suggest that the local and global intra-S checkpoints are two distinct mechanisms that can be distinguished by the signaling intensity of detecting DNA damage(Saxena and Zou, 2022). It is thought that a threshold of DNA damage must be reached to activate the global checkpoint and impact cell cycle progression during replication(Bantele et al., 2019; Shimada et al., 2002). The local checkpoint reacts to a unique replication fork stalling site, which does not impact overall cell cycle progression. The proteins involved in the global replication stress response have been identified using DNA damaging agents that lead to multiple DNA lesion formation randomly throughout the genome. Nevertheless, the mechanism and timing of the checkpoint pathway at a local scale remain largely unknown and are critical to understanding the series of processes at a single RF stall.

In the *E.coli* genome, the DNA pausing sequences called terminator (Ter) are recognized by a protein called Tus to cause a polar site-specific arrest of the RF at the end of the bacterial chromosome replication(Hiasa and Marians, 1994; Hidaka et al., 1988; Mulcair et al., 2006; Roecklein et al., 1991). The Tus-*Ter* complex leads to a temporarily locked complex on DNA that can be overcome by the fork arriving in the opposite direction, displacing Tus to terminate replication. The artificial *E.coli*-based Tus/Ter system has previously been employed in mouse embryonic stem cells to measure homologous recombination repair products and investigate the DNA repair pathway choice(Chandramouly et al., 2013; Willis et al., 2017, 2014).

In this study, we integrated a plasmid with 5 repeats of the *TerB* sequence in the non-permissive orientation, referred to as pWB15, at a unique site within chromosome 12 of the breast cancer cell line MCF7 (MCF7 5C-TerB clone). We utilized the integrated Tus-*TerB* system in human MCF7 cells to artificially generate individual RFBs and investigated the activation of the S-phase checkpoint signaling mechanism. We show that Tus/Ter creates an efficient site-specific RFB in human cells and observed local activation of ATR signaling which was responsible for the phosphorylation of DNA damage marker γH2AX at the stall sites. When a replication fork pauses at the local Tus-*TerB* block, we do not detect any alteration in global replication profiles. Our system allows us to study the ATR-checkpoint activity as a local response to a single RFB.

## Results

### Tus is found enriched at *TerB* sites integrated in human cells

The 23 bp *TerB* sequence in interaction with the Tus protein have been successfully used in yeast and mice to create artificial RFBs(Larsen et al., 2014a, 2014b; Willis et al., 2018, 2017, 2014; Willis and Scully, 2016). To study the effect of site-specific replication blocks at a genomic locus in human cells, we used a previously established MCF7 clone with a unique integrated copy of the pWB15 plasmid carrying two sets of 5 *TerB* repeat sequences in opposite, non-permissive orientations with respect to the incoming replication fork (Fig.1A, SupFig1). Whole genome sequencing of the clone was carried out to confirm the single copy integration of the plasmid at hg38 chr12:95325001. Firstly, we checked if Tus protein was able to bind *TerB sequences* in this artificial system in human cells. Chromatin-Immunoprecipitation (ChIP) followed by quantitative Polymerase Chain Reaction (qPCR) was performed using an antibody against the His-tag 24h after MCF7 5C-TerB cells were transfected with a Tus-His expression plasmid. The Tus signal is specifically enriched over 20 times compared to the vector control (VC) condition in proximity to the *TerB* sequences (PP9, 2, and 47) (Fig.1B).

**Figure 1.**
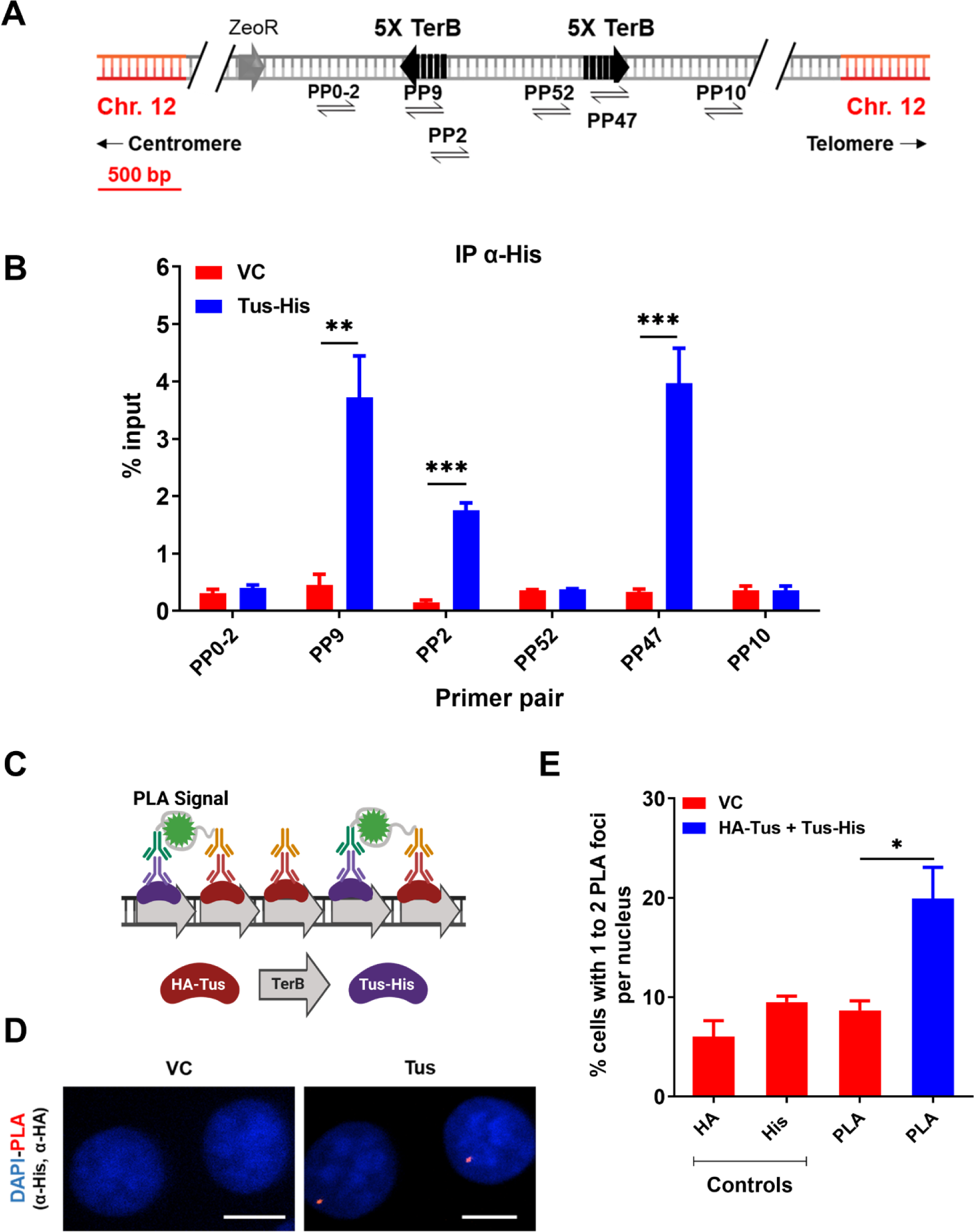
Tus is bound to *TerB* sites integrated in human cells. **(A)** Schematic of linearized plasmid (pWB15) integrated into MCF7 cells (MCF7 5C-TerB) with two cassettes containing 5 tandem *TerB* sequences in the non-permissive orientation (black arrows). Black half-arrow heads depict PCR products expected from stated primer pairs used in E. **(B)** ChIP with His antibody on MCF7 5C-TerB cells transfected with either a vector control (VC) or HA-Tus-His expression plasmids. ChIP-qPCR were conducted using the indicated primer pairs. Data show the percentage of input (n=3). **(C)** Schematic of PLA to visualize Tus bound to *TerB* sites using HA and His antibodies. **(D)** Representative images of the PLA foci across stated conditions. Bars, 10 μm **(E)** Percentage of cells with 1-2 PLA foci per nucleus. (n=3, ≥150 cells per experiment).

To corroborate this observation, we performed the proximity ligation assay (PLA) (see methods). Two tagged-Tus (HA-Tus and Tus-His) or a VC expression plasmid were simultaneously transfected and antibodies against the two tags, HA and His, were used to generate a PLA signal (Fig.1C, D). We observed 19% of the cells transfected with the two tagged Tus were harboring one or two PLA signal versus 9% of the cells transfected with the VC (Fig.1E), a significant enrichment of the PLA signal when Tus was bound to *TerB*. These results indicate a specific interaction of the Tus protein with the *TerB* sequences in human cells.

### The Tus-*TerB* barrier is an efficient block to incoming replication forks

It has previously been demonstrated that the Tus-*TerB* interaction can induce an RFB in mouse embryonic cells using a chromosomally integrated system(Willis et al., 2014). We reasoned that our integrated system in MCF7 cells would be more prone to replication fork arrests near the *TerB* sequences. We used single-molecule analysis of replicated DNA (SMARD) to investigate evidence of replication fork stalling in the genomic DNA(Norio and Schildkraut, 2001). The self-labeling SNAP-tag was used to label Tus. After induction of Myc-NLS-TUS-SNAP by using Dox for 3 days, asynchronous MCF7 5C-TerB cells were sequentially pulse-labeled using two nucleoside analogs (iododeoxyuridine (IdU) and chlorodeoxyuridine (CldU)) for 4h each (Fig.2A). The isolated DNA was digested by restriction endonuclease Sfil and the 200 kb fragment obtained was stretched on slides for staining. Two DNA FISH probes were used to detect the 200 kb segment of interest containing the *TerB* arrays: one probe of 40 kb within the chromosome 12 (hg38 chr12:95,199,827 – 95,239,226) and one probe of 7 kb made from pWB15 (Fig.2B). The direction of replication forks could be determined using the labeling patterns and its position within the locus of interest using our FISH probes. After inducing the expression of Myc-NLS-TUS-SNAP from an integrated plasmid in MCF7 5C-TerB cells (SupFig.2A, B), we analyzed its impact on the replication fork progression within the region containing *TerB* repeats (Fig.2C, D). The yellow arrows indicate the transition of labeling from IdU to CldU incorporation showing the replication fork direction. When Tus was expressed, these transition sites were found accumulated near the *TerB* regions (Fig.2D, arrows within the white oval) compared to the random signal spread on the 200 kb region when Tus is not expressed (Fig.2C). We quantified the percentage of molecules with replication forks at each 5 Kb interval in the 200 kb SfiI segment containing *TerB* sequences (Fig.2E, F) and confirmed an accumulation of forks at the *TerB* sequences only when Tus was expressed (Fig.2F, black arrow) supporting that the binding of Tus on *TerB* arrays can impair the replisome progression at genomic loci. Interestingly, expression of the Tus protein alone does not alter the direction of replication fork progression along the 200 kb SfiI segment where forks predominantly progress from the 3’ to 5’ direction in cells both with and without Tus expression as analyzed by the percentage of molecules with IdU incorporation at each 5 kb interval (SupFig.2C, D). Furthermore, no significant difference was observed in the global replication fork speed in cells with or without Tus expression (SupTable1). Together, these data indicate that when Tus bound the *TerB* sequences integrated within chromosome 12, it creates a physical barrier and an obstacle to the replication fork progression.

**Figure 2.**
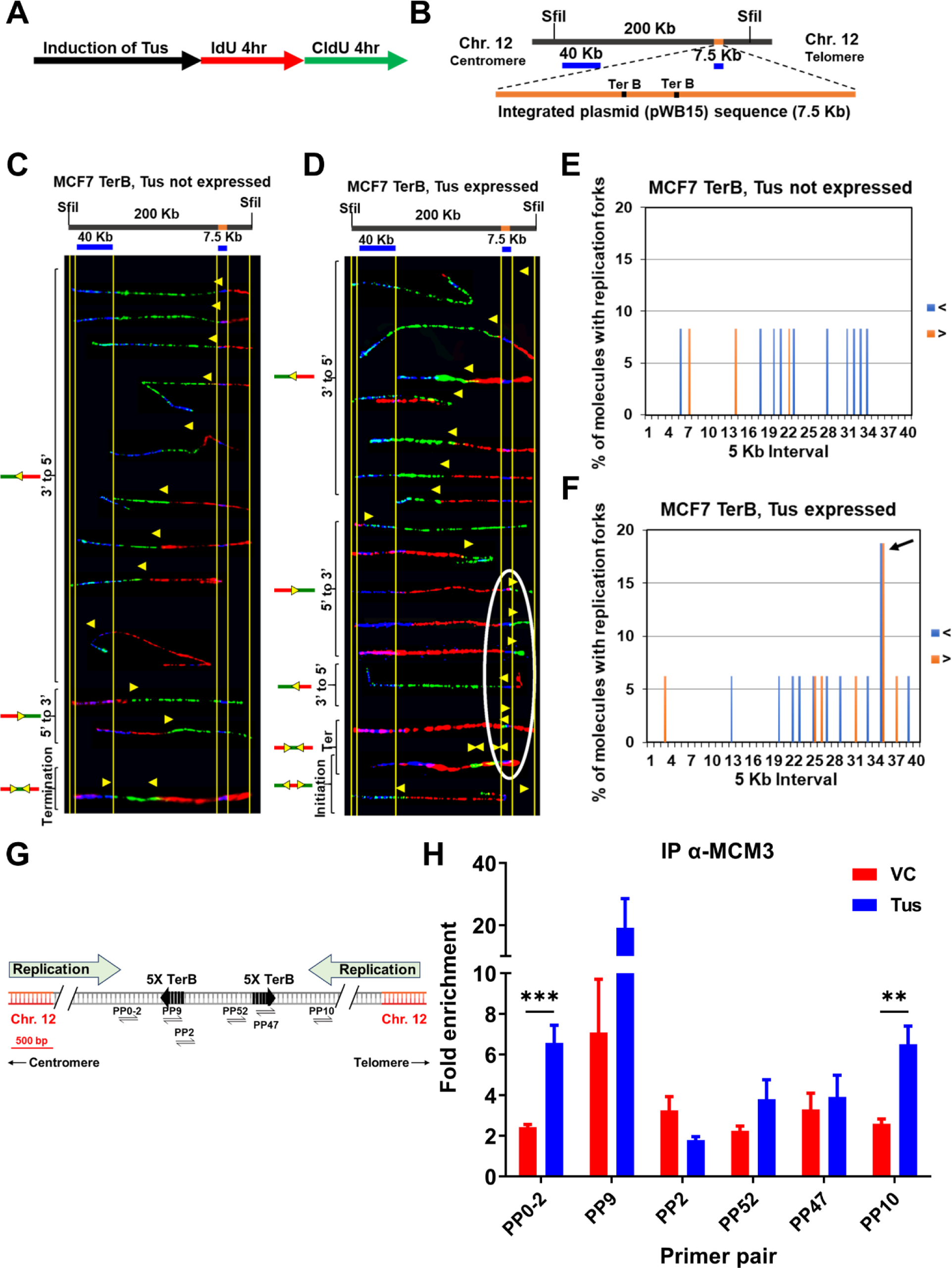
Replication fork pauses at the Ter sequence in the presence of the Tus protein. **(A)** Schematic of pulse labelling of MCF7 5C-TerB cells with Tus protein expression for 3 days. **(B)** Locus map of a 200 kb SfiI segment from MCF7 5C-TerB Chromosome 12 with the integrated plasmid (pWB15) DNA (orange). A 40 kb FISH probe made from fosmid WI2-1478M20 and a 7.5 kb FISH probe made from plasmid pWB15 are shown in blue. **(C-D)** Top: locus map of the DNA segment containing *Ter* sequence with the location of the FISH probes. Bottom: photomicrographs of labeled DNA molecules from MCF7 5C-TerB, Tus not expressed **(C)** and MCF7 5C-TerB, Tus expressed **(D)**. Yellow arrows indicate the position of replication forks at the transition of labeling from IdU to CldU incorporation showing replication fork direction. Molecules are arranged in the following order: forks progressing from 3’ to 5’, forks progressing from 5’ to 3’, termination events, and initiation events. Replication forks (Yellow Arrows) in the white oval are all at the same location (at the *TerB* sequence) on molecules from different cells indicating that replication forks are pausing at this *TerB* sequence. **(E-F)** Percentage of molecules with replication forks at each 5 Kb interval in the 200 Kb SfiI segment containing *TerB* sequence, quantified from molecules in MCF7 5C-TerB **(E)** Tus not expressed and (**F)** Tus expressed. Percentage of molecules with replication forks progressing 3’ to 5’ (< blue) and 5’ to 3’ (> orange) are shown. In the cells expressing Tus, a high percentage of molecules contain replication forks pausing in both directions in the 5 Kb interval which contains Ter sequences (black arrow **(F)**, white oval **(D)**). **(G)** Schematic similar to Fig1A with progression of the endogenous origins of replication within the chromosome 12 shown (green arrows). **(H)** ChIP using MCM3 antibody on MCF7 5C-TerB cells transfected with VC or HA-Tus-His plasmids. ChIP-qPCR were conducted using the indicated primer pairs. Data shows the fold enrichment relative to the IgG controls (n=3).

The eukaryotic replisome is a multiprotein apparatus consisting of DNA helicases, DNA polymerases and accessory proteins to duplicate the genome. We hypothesized that an RFB would induce an accumulation of these proteins on chromatin around *TerB* sequences. To test our hypothesis, we performed ChIP using an antibody against mini-chromosome maintenance protein 3 (MCM3), a CMG core-complex protein, in the absence or presence of Tus. When Tus was expressed, MCM3 was 3-fold more enriched upstream of the *TerB* arrays at the PP0-2 and PP10 sites compared to the surrounding primer pairs (Fig.2G, H). Additionally, we find that there is an enrichment of the DNA repair scaffold protein FANCM at the *TerB* sites mirroring the MCM3 enrichment (SupFig.3), indicative of the stalled replication forks as previously described(Collis et al., 2008; Nandi and Whitby, 2012; Panday et al., 2021; Xue et al., 2015). Together, these results show that Tus expression induces a RFB at the *TerB* arrays impairing the replisome progression during S-phase.

**Figure 3.**
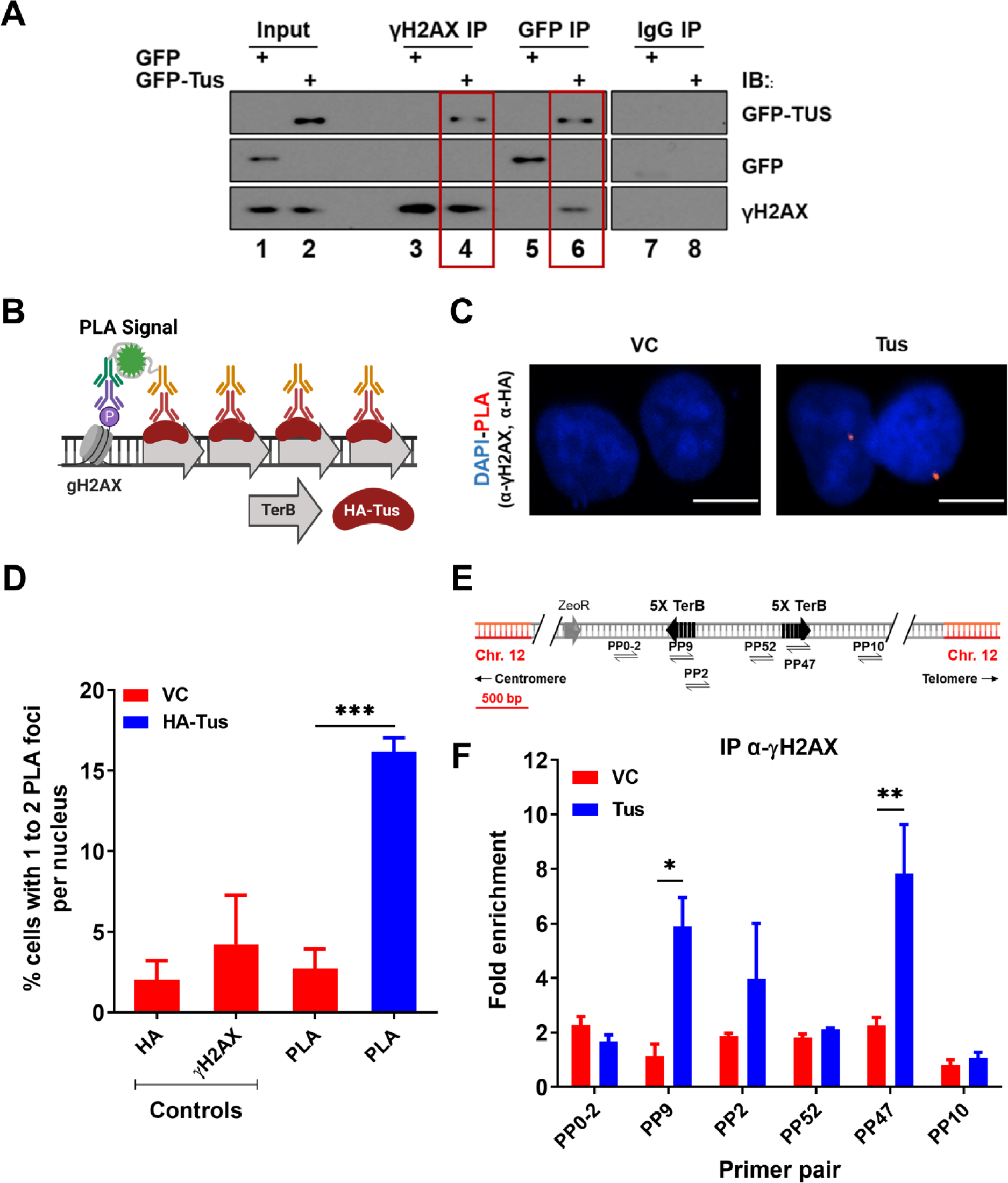
γH2AX is enriched at *TerB* sites after Tus expression. **(A)** MCF7 5C-TerB cells transfected with GFP or GFP-Tus, γH2AX and IgG antibodies were used to immuno-precipitate proteins and analysed by immunoblotting with indicated antibodies. **(B)** Schematic of PLA to visualize HA-Tus bound to *TerB* in the proximity of γH2AX sites using HA and γH2AX antibodies **(C)** Representative images of the PLA foci across stated conditions. Bars, 10 μm **(D)** Percentage of cells with 1-2 PLA foci per nucleus. (n=3, ≥150 cells per experiment). **(E)** Schematic of MCF7 5C-TerB depicting the positions of the primer pairs with respect to the integrated *TerB* locus. **(F)** γH2AX levels along the integrated *TerB* plasmid were analyzed by ChIP-qPCR in MCF7 5C-TerB cells transfected with VC or Tus expression plasmids using the indicated primer pairs. Data shows the fold enrichment relative to IgG controls (n=3).

### γH2AX is enriched at *TerB* sites after Tus expression before fork collapse

One of the earliest responses to replication stress is the phosphorylation of the histone variant H2AX on serine 139 (γH2AX) by members of the phosphoinositide 3-kinase (PI3K)-like family (PIKK)(Ward and Chen, 2001). As we demonstrated that *TerB* arrays can block incoming replication forks, we asked whether we could see a γH2AX signal enrichment due to fork arrest. Using a co-immunoprecipitation assay with the chromatin fraction of cross-linked-cells expressing either GFP tag or GFP-Tus, we found that GFP-Tus and γH2AX were immunoprecipitated together using GFP or γH2AX antibodies (Fig.3A).

To gain insight into the γH2AX signal at the RFB induced by the Tus-*TerB* interaction, cells were transfected with either VC or HA-Tus expression plasmids, and PLA was performed using antibodies against HA and γH2AX (Fig.3B). We found that 16% of the HA-Tus transfected cells were harboring at least one PLA signal versus 3% of the VC transfected cells (Fig.3C, D).

To better characterize the γH2AX enrichment around *TerB* sites, we performed ChIP-qPCR assays using a γH2AX antibody 24h after Tus or VC expression. We observed an enrichment of γH2AX, strictly co-localizing with the Tus-Ter interaction sites (PP9 and PP47), suggesting a local role of γH2AX in response to stalled RFs (Fig.3E, F). This is in stark contrast to the observed spreading of the γH2AX signal after a site-directed DSB(Berkovich et al., 2007; Chailleux et al., 2014; Clouaire et al., 2018; Savic et al., 2009). To corroborate this observation in our integrated cassette, we generated a site-specific DSB by expressing Cas9 targeted to the integrated cassette (SupFig.4A-C). We assessed γH2AX using the ChIP-qPCR assay 24h after transfection and noticed enrichment at the distal primer pairs (PP0-2 and PP10), contrasting with the very tight local signal observed at the Tus-*TerB* RFB (SupFig.4D). However, we did not find any significant γH2AX signal between VC or Tus expression (SupFig 5A-B) supporting the lack of global response. This suggests that the RFB generated by the interaction between Tus and the *TerB* arrays activates a stress response that stimulates the phosphorylation of H2AX, concentrated around the fork block.

**Figure 4.**
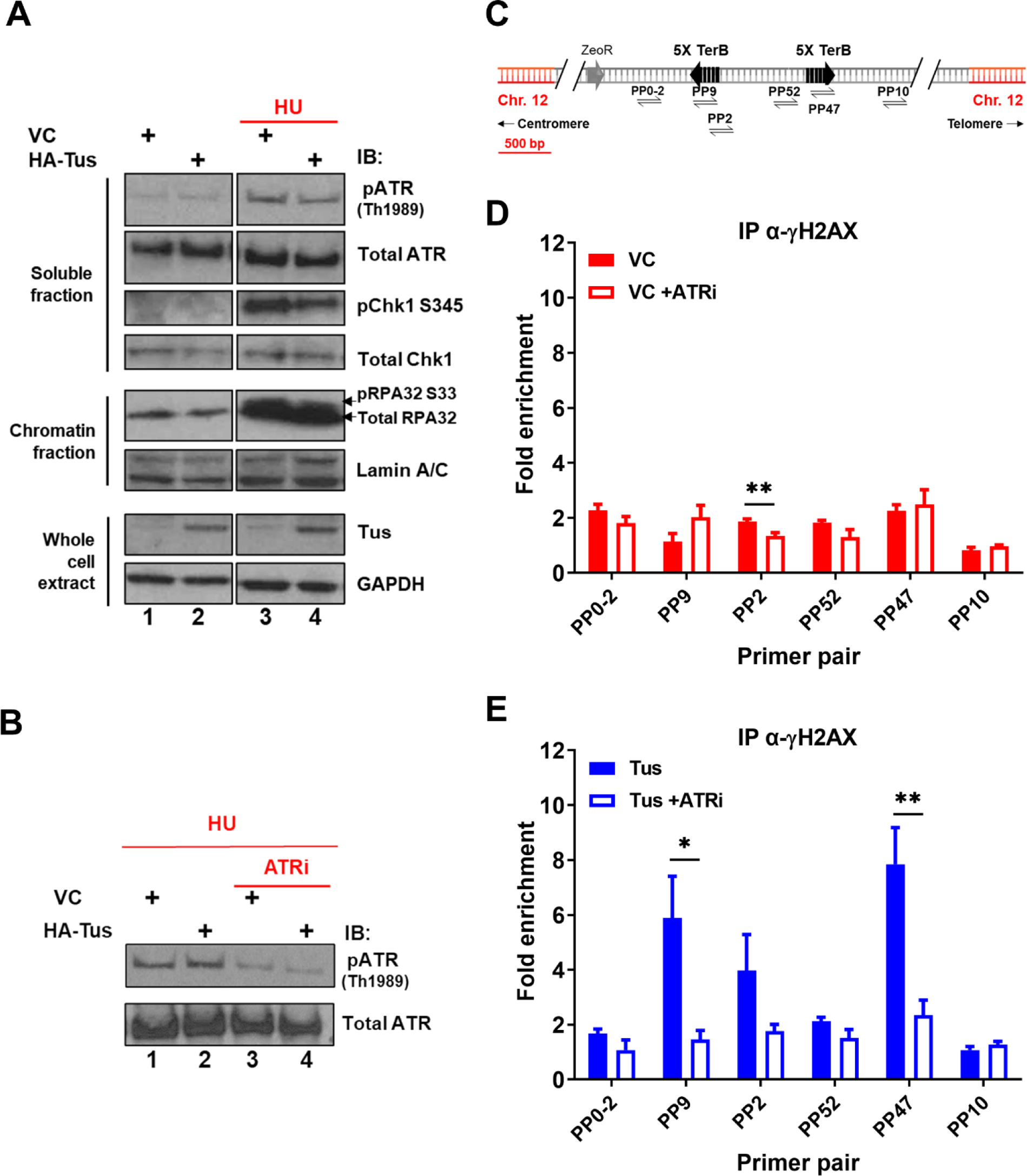
γH2AX phosphorylation at Tus-*TerB* is ATR-dependent. **(A)** MCF7 5C-TerB cells transfected with VC or HA-Tus were treated for 4hrs with 2mM HU before lysis and fractionation (whole cell extract (WCE), soluble fraction and chromatin fraction). Total and phosphorylated protein levels were examined by immunoblotting as indicated. Tus expression was confirmed in WCE. **(B)** Depletion of pATR Th1989 in MCF7 5C-TerB cells treated with 2mM HU +/- 40 nM of ATR inhibitor was examined with immunoblotting. **(C)** Schematic of MCF7 5C-TerB depicting the positions of the primer pairs with respect to the integrated *TerB* locus. **(D-E)** γH2AX levels were analyzed by ChIP-qPCR in MCF7 5C-TerB cells transfected with **(D)** VC expression plasmid **(E)** Tus expression plasmid, +/- 4hrs treatment with ATRi with indicated primer pairs. Data shows the fold enrichment over the IgG controls (n=3).

**Figure 5.**
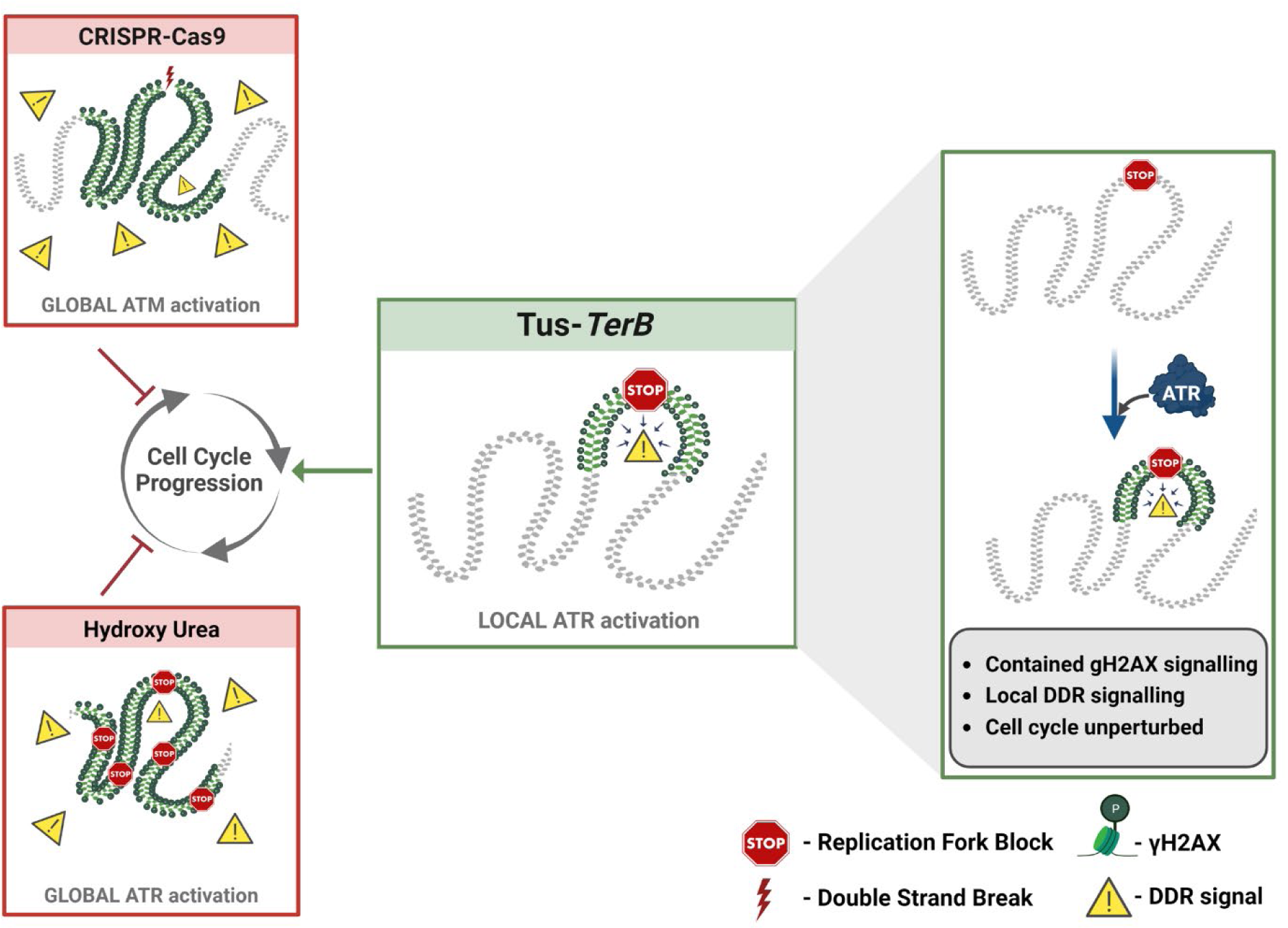
A local ATR-dependent checkpoint is activated by the Tus-*TerB* RFB. A model depicting the cellular response to a site-specific DSB using CRISPR-Cas9 and a global replication stress with hydroxyurea, both of which lead to a DNA damage response (DDR, yellow triangles), increased gH2AX (green) levels globally in the cell and, if left unresolved, can alter cell cycle progression. In contrast, the site-specific replication fork block, Tus/*TerB*, elicits a local ATR dependent DNA damage response which is responsible for the phosphorylation and accumulation of γH2AX at the stalled site. This local signaling does not affect the progression of the cell cycle is not altered during the local checkpoint response.

### H2AX is phosphorylated in an ATR-dependent manner without a global activation of the intra-S checkpoint

ATR is known to phosphorylate H2AX in response to replication stress and plays a pivotal role in the surveillance of DNA replication(Ward and Chen, 2001). To understand if the global ATR-dependent checkpoint was activated in response to the Tus-*TerB* RFB, we analyzed the levels of phospho-proteins involved in the intra-S checkpoint activation pathway (RPA, CHK1, and ATR) by immunoblotting fractionated cell extracts 24h after VC or Tus expression. Cells treated with 2mM hydroxyurea (HU) for 4h demonstrated an increase in phospho-protein levels, acting as a control for checkpoint activation (Fig 4.A, lanes 3 and 4). Tus expression alone did not activate the global ATR-dependent checkpoint as the level of phospho-protein remained the same as the VC control (Fig 4.A, lanes 1 and 2). We did not find any observable difference in cell cycle progression between VC or Tus expression alone (SupFig6 A-B) supporting the lack of global response.

We hypothesized that if the local γH2AX enrichment at *TerB* sites was ATR-dependent, a reduction of ATR activity would lead to a decreased γH2AX signal. To validate the ATR inhibitor activity (ATRi), VE-822(Fokas et al., 2012), we performed immunoblotting using total protein extracts of cells treated with 2mM HU and compared the level of phospho-ATR TH1989 with or without an ATR inhibitor. A 4h treatment with ATRi reduced the level of phospho-ATR TH1989, reflecting a decrease in ATR activity (Fig.4B). ChIP-qPCR assays were performed using a γH2AX antibody 24h after VC or Tus expression and cells were harvested after a 4h ATRi treatment. Consistent with our hypothesis, the γH2AX enrichment in the vicinity of *TerB* sites after Tus expression was remarkably decreased upon ATRi treatment (Fig.4C-E PP9 and PP47), implying that the Tus-*TerB* RFB activates a localized stress response that stimulated the local phosphorylation of H2AX. Together, these data suggest a local activation of the intra-S checkpoint via the ATR kinase and rapid phosphorylation of H2AX, which could mediate the recruitment of repair factors near the damage site.

## Discussion

Replication stress occurs during the S-phase of the cell cycle, when the replication machinery encounters DNA lesions that cause stalling of replicative polymerases and can be a significant cause of genomic instability. If the stalled forks cannot be processed, they can result in DNA breakage, mutations, and chromosomal rearrangements leading to the development of many different human cancers(Gaillard et al., 2015; Tubbs and Nussenzweig, 2017). In this study, we used the reconstituted *E.coli*-derived protein-DNA barrier, Tus-*TerB,* to study the replication stress response at a single RFB. We have shown that the Tus protein efficiently binds at the *TerB* sequences using ChIP-qPCR and PLA in MCF7 cells (Fig.1). We confirmed that this integrated system was able to cause site-specific RFB at *TerB* sequences when Tus was introduced with the use of the SMARD technique and observed the accumulation of replication fork protein MCM3 upstream of *TerB* sequences (Fig.2). Our results indicate that the integrated Tus-*TerB* system acts as an efficient site-specific replication fork barrier, providing valuable insights into the local processing of a stalled single fork in human cells and its difference from global replication stress (Fig.5).

We show that the site-specific replication stress leads to the accumulation of γH2AX at the site of fork block when Tus was expressed by co-immunoprecipitation of Tus and γH2AX (Fig.3A). Additionally, we confirmed that γH2AX is localized at the site of Tus-*TerB* using PLA and ChIP-qPCR assays (Fig.3D,F). We suggest that the γH2AX signal is being constrained to a region of less than a kilobase from the *TerB* sites by tortional stress generated when the replication fork encounters the barrier and is not influenced by well-defined topological domains that are one measure of the functional units of the genome(Dixon et al., 2012; Rao et al., 2014). Furthermore, there was no observable difference in global replication profiles or the activation of global checkpoint markers such as pRPA, pCHK1, and pATR, with or without Tus protein expression, contrasting with the differences observed in the presence and absence of HU-induced replication stress (Fig.4A, SupTable1 & SupFig.5,6). We conclude that Tus-*TerB*-induced replication fork stall does not activate global ATR-dependent S-phase checkpoint signaling, suggesting that a local response is occurring at the stalled replication fork. Interestingly, upon ATR inhibition, we observed that the enrichment of phosphorylated H2AX at *TerB* sites was significantly reduced when Tus was expressed, which indicates local ATR signaling is responsible for the phosphorylation of H2AX at the stalled site (Fig.4C,E). A similar localized response had previously been observed in yeast, with a site-specific DSB in G1 cells, where there was no activation of the global checkpoint signaling, but γH2AX signaling was observed (Janke et al., 2010). This localized ATR-dependent γH2AX differs from the well-observed ATM-dependent spreading of γH2AX signal after a site-directed double-strand break(Berkovich et al., 2007; Chailleux et al., 2014; Clouaire et al., 2018; Savic et al., 2009) and suggests a distinct local intra-S phase checkpoint is being activated in response to the single RFB.

There are additional mechanisms to be understood in this newly described local replication stress response, which includes whether the activation of ATR is dependent on ATRIP bound to ssDNA (Ball et al., 2005; Zou and Elledge, 2003). In the Tus-*TerB* system, there is no known dissociation of the replicative helicase from the polymerase, which is responsible for the ssDNA signal. Therefore, just like ATM can be activated independently of the MRE11 complex(Bakkenist and Kastan, 2003), we expect that ATR is activated independently of ATRIP, likely due to local changes in chromatin that will be investigated in future studies.

During extensive DNA damage, the genome-wide checkpoint response activates ATR/CHK1 globally, which leads to the slowing of all replication forks in the cell, inhibition of cell cycle progression, and suppression of late origin firing; to provide sufficient time for DNA repair. It has been proposed that a local checkpoint response occurs when one or a few replication forks encounter a DNA lesion and activates ATR/CHK1 signaling at local sites of fork stalling. The local ATR response has been hypothesized to be transient, in which the fork moves slowly only at stalled sites to promote fork stabilization, restart the stalled fork, and suppress recombination without triggering the global checkpoint response(Iyer and Rhind, 2017, 2013; Kaufmann et al., 1980; Merrick et al., 2004; Saxena and Zou, 2022; Willis and Rhind, 2009b; Zeman and Cimprich, 2013).

We predict that the local ATR-dependent checkpoint signaling can result in a rapid and controlled response to allow RFB resolution, through the recruitment of repair factors to the damaged site. The factors might include those involved in fork stabilization, fork reversal, fork cleavage, and homology-dependent replication restart(Saxena and Zou, 2022). The Tus-*TerB* system has previously been employed in mouse embryonic stem cells to measure and investigate the DNA repair pathway choice(Chandramouly et al., 2013; Panday et al., 2021; Willis et al., 2017, 2014). This included both error-free and error-prone homologous recombination induced by a mammalian chromosomal RFB, as well as identifying the role of the structure-specific endonuclease complex SLX4-XPF and FANCM in the repair of microhomology-mediated tandem duplications that occur at replication arrest(Elango et al., 2022; Nandi and Whitby, 2012; Willis et al., 2017). The Tus-*TerB* block is robust and is not dissociated by the action of DNA helicases such as PIF1, corroborated in previous yeast studies that have demonstrated that the “sweepase” RRM3 is unable to resolve this RFB(Larsen et al., 2014b). Therefore, it is likely that the RFB is resolved either by incoming replication forks leading to termination at the *TerB* site or by fork cleavage(Larsen et al., 2014b; Willis et al., 2017, 2014). Fork reversal at the Tus-*TerB* site has been observed in the yeast system(Larsen et al., 2014b; Marie and Symington, 2022), but not for the mammalian systems, and it is unlikely that shifting the position of the RFB would make it easier to resolve. Therefore, fork cleavage is the most likely mechanism for resolving the RFB.

The upstream pathway of activation of the DNA damage response, as a result of a single RFB, is not completely understood. Previous work has shown that FANCM acts as a scaffolding protein for recruitment of different repair protein complexes involved in the stalled replication fork rescue(Panday et al., 2021; Willis et al., 2017). We observed that the FANCM protein is enriched at the site of the RFB induced by Tus-*TerB* (SupFig.3), confirming a similar phenomenon observed in mouse cells. However, it is yet to be determined if FANCM recruitment to the RFB precedes local H2AX phosphorylation by ATR, or if FANCM is recruited as a consequence of the γH2AX signal to help elicit the downstream DNA damage response (DDR). We also anticipate that the protein complex, 9-1-1 (RAD9A, HUS1, RAD1), and the subsequently recruited TOPBP1, may also play a role at this local ATR-dependent intra-S checkpoint(Helt et al., 2005; Parrilla-Castellar et al., 2004).

While the Tus-*TerB* system is an important tool in deciphering signaling at individual replication forks, it will be important to test if the local-S phase model remains supported when exploring endogenous lesions that occur when replication forks encounter secondary structures such as R-loops, G-quadruplexes and common fragile sites characterized by trinucleotide repeat expansion [(CAG)_n_/(CTG)_n_] (Brickner et al., 2022; Bryan, 2019; Kim et al., 2016). Here the mechanisms surrounding the identification and resolution of the RFB may differ. Overall, our results indicate that Tus-*TerB* system acts as an efficient RFB and activates local ATR checkpoint signaling at the stall site, leading to phosphorylation and accumulation of the DNA damage sensor protein γH2AX, which is dependent on the ATR kinase. The local γH2AX accumulation at the stalled region would lead to the recruitment of DNA repair factors for resolution of the fork (Fig.5). Together, our findings reveal the signaling mechanism of the Tus/*TerB* induced replication block and showed that the site-specific fork block is dependent on the local ATR S-phase checkpoint signaling.

## Methods

### Cell culture and transfection conditions

MCF7 5C-TerB cells were grown at 37°C and 5% CO_2_ in complete DMEM high glucose supplemented with 10% FBS, 2 mM Glutamax, 20 mM HEPES, 100 I.U./ml Penicillin, and 100 μg/ml Streptomycin supplemented with 400 µg/ml zeocin.

For PLA, cells were seeded at 1X10^5^ in a 12-well cell culture plate. For T7 assay, cells were seeded at 2.5X10^5^ in a 6-well cell culture plate. For ChIP, IP and cellular fractionation, cells were seeded at 2.5X10^5^ in a 10 cm cell culture dish.

MCF7 5C-TerB cells were seeded and transfected the day after with 10 μg of pCMV3xnls Tus or pCMV3xnls using the Mirus Bio™ TransIT™-LT1 reagents for 24h more. For the generation of DSB, cells were transfected 6h prior harvesting with 120 pmol of the sgRNA TerB1 and 120pmol of the Cas9 protein (obtained from QB3 Macrolab, Berkeley) using the Lipofectamine CRISPRMAX™ Cas9 transfection reagents.

### Generation of stable cell lines

PvuI linearized pWB15 was transfected into MCF7 cells to generate the stably integrated clone, MCF7 5C-TerB (Supplementary Figure 1). 800 μg/ml of Zeocin was used to select positively integrated cells and monoclonal colonies were isolated and subsequently maintained in 400 μg/ml Zeocin. Single integrant validation was performed by whole genome sequencing of the clone.

Stable MCF7 5C-TerB cells inducible expressing Myc-NLS-TUS-SNAP were generated by lentiviral transduction. Cells were infected with lentiviral particles containing pInducer10L or pInd Tus-SNAP and single-cell clonal colonies were selected in complete DMEM containing 1 µg/mL puromycin and 400 µg/ml zeocin. Myc-NLS-TUS-SNAP protein was expressed in stable lines by induction with 1 µg/ml doxycycline for 3 days.

### Plasmid construction

The doxycycline-inducible lentiviral plasmid used to express N-terminally myc-tagged, C-terminally SNAP-tagged nuclear localized, human codon-optimized wild-type Tus (Myc-NLS-TUS-SNAP) was generated as follows. The N-terminally myc epitope-tagged, nuclear-localized, codon-optimized wild-type Tus cDNA sequence was PCR amplified from pcDNA3-β-MYC-NLS-Tus using primers Forward 5’-AGTCGGTACCGAATTCGCCACCATGGAACAAAAGCTG-3’ and Reverse 5’-AGTCGGCGGCCGCGCCGCTACCGTCAGCCACGTACAGGTGCA and the SNAP-tag cDNA sequence PCR amplified using primers Forward 5’-AGTCGCGGCCGCCGGCCACATGGACAAAGACTGCGAAATGAAGC-3’ and Reverse 5’-ACTGCTCGAGTCAACCCAGCCCAGGCTTGC-3’. The Tus and SNAP PCR amplified fragments were digested with KpnI and NotI, and NotI and XhoI, respectively, and inserted into the inducible expression lentiviral vector pInducer10L(Drosopoulos et al., 2020) directly downstream of the doxycycline-inducible promoter, to generate pInd Tus-SNAP. This construct was sequenced to confirm that unintended mutations were not introduced during PCR and cloning.

### Immunoblotting

Cells were harvested by trypsinization, suspended in complete DMEM, washed with PBS, then pelleted and flash frozen in liquid N_2_ and stored at -80°C. For SDS-PAGE, pellets were thawed on ice and lysed by resuspending in Laemmli Buffer (60 mM Tris-HCl pH 6.8, 400 mM 2-mercaptoethanol, 2% SDS, 10% glycerol, 0.01% bromophenol blue) to a final concentration of 10^6^ cells/ml. Lysates were denatured (5 min at100°C) and passed through a 25-gauge needle (5x) then spun for 2 min at full speed in a microfuge. Aliquots of lysate corresponding to 10^5^ cells were resolved on 4-15% gradient SDS-PAGE gels, proteins transferred to nitrocellulose membrane and blocked in PBS with 5% Blotting-grade Blocker and 0.1% Tween 20. Membranes were then incubated with primary antibodies diluted in PBS with 5% Blotting-grade Blocker. Primary antibodies used were anti-Myc tag and anti-actin. Following incubation with primary antibodies, membranes were washed with PBS + 0.1% Tween 20. Membranes were then incubated with fluorescently labeled Goat Anti-Mouse IRDye 680LT and Goat Anti-Rabbit IRDye800CW secondary antibodies and then washed in PBS + 0.1% Tween 20. Immunoblots were imaged on an Odyssey Lc Infrared scanner.

### Single Molecule Analysis of Replicated DNA (SMARD) assay

SMARD was performed essentially as previously described(Drosopoulos et al., 2012; Norio and Schildkraut, 2001; Twayana et al., 2021). Exponentially growing cells in complete DMEM with 1 µg/ml doxycycline were sequentially pulse-labeled with IdU (4 h) followed by CldU (4 h) each to a final concentration of 30 μΜ. Following pulsing, cells were suspended in PBS at a concentration of 3 × 10^7^ cells per ml. An equal volume of 1% molten TopVision low melting point agarose) in PBS was added. The resulting cell suspension in 0.5% TopVision agarose was poured into wells of a chilled plastic mold to make plugs of size 0.5 cm × 0.2 cm × 0.9 cm, each containing 10^6^ cells. Cells in the plug were lysed at 50°C in a buffer containing 1% n-lauroylsarcosine, 0.5 M EDTA, and 0.2 mg/ml proteinase K. Plugs were rinsed in TE and washed with 200 μM phenylmethanesulfonyl fluoride (PMSF). Genomic DNA in the plugs was digested with *SfiI* overnight at 37°C. Digested genomic DNA was cast into 0.7% SeaPlaque GTG agarose gel and DNA was separated by pulsed field gel electrophoresis (PFGE) using a CHEF-DRII system (Bio-Rad). The 200 kb DNA segment containing *TerB* sequences within the gel was located by performing Southern hybridization on a portion of the gel using a specific probe made from the pWB15 plasmid. This identified the pulse field gel slice containing *TerB* sequences which was excised and melted (20 min at about 72°C). The DNA in the gel solution was stretched on microscope slides coated with 3-aminopropyltriethoxysilane, denatured with sodium hydroxide in ethanol, fixed with glutaraldehyde and hybridized overnight with biotinylated DNA FISH probes at 37°C in a humidified chamber. Biotinylated DNA FISH probes were made from the fosmid WI2-1478M20 and from the plasmid pWB15. Following hybridization, slides were blocked with 3% BSA for at least 20 min and incubated with the avidin Alexa Fluor 350. This was followed by two rounds of incubation with the biotinylated anti-avidin D for 20 min followed by the avidin Alexa Fluor 350 for 20 min. Slides were then incubated with an antibody specific for IdU; a mouse anti-BrdU antibody, an antibody specific for CldU; a rat monoclonal anti-BrdU antibody, and the biotinylated anti-avidin D for one hr. This was followed by incubation with secondary antibodies: Alexa Fluor 568 goat anti-mouse IgG (H+L), Alexa Fluor 488 goat anti-rat IgG (H+L), and the avidin Alexa Fluor 350 for one hour. Slides were rinsed in PBS with 0.03% IGEPAL CA-630 after each incubation. After the last rinse in PBS/CA-630, coverslips were mounted on the slides with Prolong gold antifade reagent. A Zeiss fluorescent microscope and IP Lab software (Scanalytics) were used to capture images of IdU/CldU incorporated DNA molecules. Images were processed using Adobe Photoshop and aligned (using Adobe Illustrator) based on the positions of FISH signals that identify the 200 kb segments that contain the ter sequences.

### Proximity Ligation Assay (PLA)

MCF7 5C-TerB cells were seeded on poly-L-lysine coated coverslips and transfected the day after as described before. 24h hours after transfection, cells were washed with PBS, and fixed and permeabilized with 4% paraformaldehyde containing 0.2% Triton X-100 for 20min on ice and blocked with PBS-BSA 3% overnight at 4. Coverslips were incubated with primary antibodies (see Table 1 for dilution) for 1h at RT. Proximity ligation was performed using the Duolink® In Situ Red Starter Kit Mouse/Rabbit (Sigma-Aldrich) according to the manufacturer’s protocol. The oligonucleotides and antibody-nucleic acid conjugates used were those provided in the Sigma-Aldrich PLA kit. Images were quantified by counting the number of foci per nucleus using Nikon software.

### Immunoprecipitation

For immunoprecipitation, MCF7 5C-TerB cells were pre-extracted using CSK100 buffer (100 mM NaCl, 300mM sucrose, 3 mM MgCl2, 10 mM PIPES pH 6.8, 1 mM EGTA, 0.2%

Triton X-100, anti-protease and anti-phosphatase) for 5 min on ice, fixed in 1% formaldehyde for 10 min on ice and a 1% glycine solution was used to stop the reaction. After scraping the cells in ice-cold PBS and centrifugation, pellets were lysed in SDS lysis buffer (50 mM Tris-HCl pH 7.5, 150 mM NaCl, 0.1% SDS, anti-proteases and anti-phosphatases) for 10 min on ice and sheared for 3 min. Samples were cleared by centrifugation for 5 min at 4°C. Immunoprecipitations were performed on 10-fold diluted lysates in dilution buffer (50 mM Tris-HCl pH 7.5, 150 mM NaCl,5 mM EDTA, 0.2% Triton X-100, anti-proteases and anti-phosphatases). with antibodies against GFP, γH2AX or IgG overnight at 4°C on a wheel. Beads were extensively washed in the dilution buffer and denatured in 2X Laemmli buffer.

Proteins were separated on 4-12% acrylamide SDS-PAGE, transferred on Nitrocellulose membrane and detected with the indicated antibodies described in the table and ECL reagents.

### Cellular fractionation

For the cellular fractionation, MCF7 5C-TerB cells were scraped in PBS, divided in 2 different tubes (1/3 of the volume for the whole cell extract (WCE) and the remaining 2/3 for the fractionation) and centrifuged to keep the pellets.

For the WCE, cells were lysed in 1 volume of lysis buffer (50 mM Tris Ph 7.5, 20 Mm NaCl, 1 Mm MgCl2, 0.1% SDS, anti-protease and anti-phosphatase) for 10 min at RT on a wheel and denaturated in 2X Laemmli buffer.

For the fractionation, MCF7 5C-TerB cells were pre-extracted in 2 volumes of CSK100 (100 mM NaCl, 300mM sucrose, 3 mM MgCl2, 10 mM pipes pH 6.8, 1 mM EGTA, 0.2% Triton X-100, anti-protease and anti-phosphatase) for 15 min on ice, and centrifuged. The supernatant (SN) representing the soluble fraction was kept in a new tube (soluble fraction). The pellet was washed with CSK50 (50 mM NaCl, 300mM sucrose, 3 mM MgCl2, 10 mM pipes pH 6.8, 1 mM EGTA, 0.2% Triton X-100, anti-protease and anti-phosphatase and resuspended in 2 volumes of CSK50 containing benzonase for 10 min on a rotating wheel, and the SN was kept after centrifugation (chromatin fraction). All the fractions were denaturated in 2X Laemmli buffer.

Proteins were separated on 4-12% acrylamide SDS-PAGE, transferred on Nitrocellulose membrane and detected with the indicated antibodies described in the table and ECL reagents.

### Chromatin Immunoprecipitation Assay

ChIPs were performed using the ChIP-IT express according to manufacturer’s instructions. Cells were crosslinked, washed, harvested by scraping and lysed according to the kit instructions. Chromatin was sonicated to 200-1500 bp fragments and DNA concentration determined. Equal amount up to 15 ug of DNA was then used for each immunoprecipitation. Antibodies used were listed on the table below. DNA fragments were eluted, purified, and analyzed by SYBR Green real-time PCR. Sequences of primers used for qPCR are given in the table. Experiments were repeated at least three times and each real time qPCR reaction was performed in duplicate.

### Detection of DSBs using the T7 endonuclease assay

MCF7 5C-TerB cells were seeded and transfected the day after with 120 pmol of Cas9 protein and 120 pmol of the sgRNA TerB1 using the Lipofectamine CRISPRMAX™ Cas9 transfection reagents (previously described). 24 hours post-transfection, cells were lysed using a lysis buffer (100 mM NaCl, 10 mM Tris-HCl pH 8, 25 mM EDTA pH 8, 0,5% SDS) and 50 ug of Proteinase K overnight at 50°C with shaking. After addition of NaCl and mix for 1 min, the supernatant was put in a new tube and an ethanol precipitation was performed. A PCR to amplify the 900 bp region surrounding the sgRNA TerB1 site was conducted using PP9F and R. Heteroduplex were formed by heating and cooling down the samples and a T7 endonuclease assay digestion was performed prior electrophoresis for detection.

### Cell Cycle progression

MCF7 5C-TerB cells were transfected as described before. Control cells were treated with 2mM HU for 4 hours before harvesting. Cells were washed with PBS and fixed in 1 ml cold 70% ethanol for 30 minutes. Cells were pelleted and washed with PBS. 100 µg/mL RNase A was added 50 µg/mL PI solution was added directly to the pellet, mixed well and incubated for 30 mins at room temperature in the dark. 50,000 cells per condition were analyzed by flow cytometry.

### Immunofluorescence (γH2AX)

Cells were fixed with 4% PFA/PBS for 20 min, permeabilized with 0.5% Triton-X 100 for 10 min, washed 3 times in 1X PBS, and blocked in 10% goat serum overnight at 4°C. The primary antibody (1:500) was incubated for 2 hr at room temperature. Cells were then washed 3 times in 1X PBS. The secondary antibody (1:1500) was incubated for 1 hr at room temperature in dark. Cells were then washed again 3 times in 1X PBS. Stained cells were mounted with mounting medium containing DAPI. Slides were imaged at 60X (immersion oil) using Nikon A1 spinning disk confocal microscope.

### Image Analysis

For PLA and gH2AX experiments, slides were imaged at 60X (immersion oil) with Nikon spinning disk confocal microscope. PLA foci per nucleus and gH2AX foci per nucleus were calculated using Nikon Elements AR Analysis Explorer (version 5.21.03), where DAPI was used as a mask for the nucleus. The number of PLA foci per nucleus were quantified to get the percentage of cells with 1 or 2 foci indicating a positive signal at the site-specific replication block. The number of gH2AX foci was counted for each DAPI to obtain the average number of gH2AX foci in each condition.

### NGS sequencing of the MCF7 5C-TerB clone

Genomic DNA was extracted from the cells and submitted to Novogene for sequencing. The samples were processed and whole genome sequencing was performed using their hWGS service pipeline.

### Data analysis and Statistics

Statistical analysis for the various experiments was performed using GraphPad Prism version 9.4.0. Results are presented as mean ± standard error of the mean (SEM). A p value of <0.05 by Student’s t-test was considered statistically significant. ns: non-significant, * Indicates p<0.05, **: P<0.005, ***: P<0.001.

## Data availability

The list of antibodies, plasmids and oligonucleotides used in this study are provided in Supplementary Table 2. WGS sequencing files generated in this study have been deposited at the Gene Expression Omnibus (GEO) database under accession number PRJNA868342 and are publicly available as of the date of publication. Any additional information required to reanalyze the data reported in this paper is available from the lead contact upon request.

## Acknowledgments

We are indebted to Ino de Bruijn, Jorge S. Reis-Filho, and Yingjie Zhu for help with the genomics data. We thank members of the Powell and Schildkraut lab for their comments on the manuscript. This work was supported in part by NIH Grants R01-CA187069 and P50-CA247749 (to S.N.P.); 5R01-GM045751 and R01-CA085344 (to C.L.S.); National Cancer Institute Cancer Center Support Grants, P30-CA008748 at MSK, and P30-CA013330 for use of a core facility at Albert Einstein COM. S.T. was supported by NIH Training Grant T-32 NIH 5T32AG023475. Figures were created with the aid of BioRender.

## Author Contributions

Conceptualization, S.N.P., S.A.S. and C.L.S.; Methodology, S.N.P., S.A.S., M.J., H.E.G., A.S., C.T. and C.L.S; Validation, S.A.S, M.J. and C.L.S.; Formal Analysis, S.A.S, M.J., H.E.G., A.S., C.T. and C.L.S.; Investigation, S.A.S, M.J., S.T., H.E.G., A.S., C.T. and S.T.K. ; Resources, S.N.P. and C.L.S ; Writing – Original Draft, S.A.S., M.J., H.E.G., A.S. and S.N.P.; Writing – Review & Editing, C.L.S and S.T.K.; Visualization, S.A.S., M.J., H.E.G. and A.S.; Supervision, S.N.P. and C.L.S.; Project Administration, S.N.P. and C.L.S.; Funding Acquisition, S.N.P. and C.L.S.

## Supplementary Figures

**Supplementary Figure 1.**
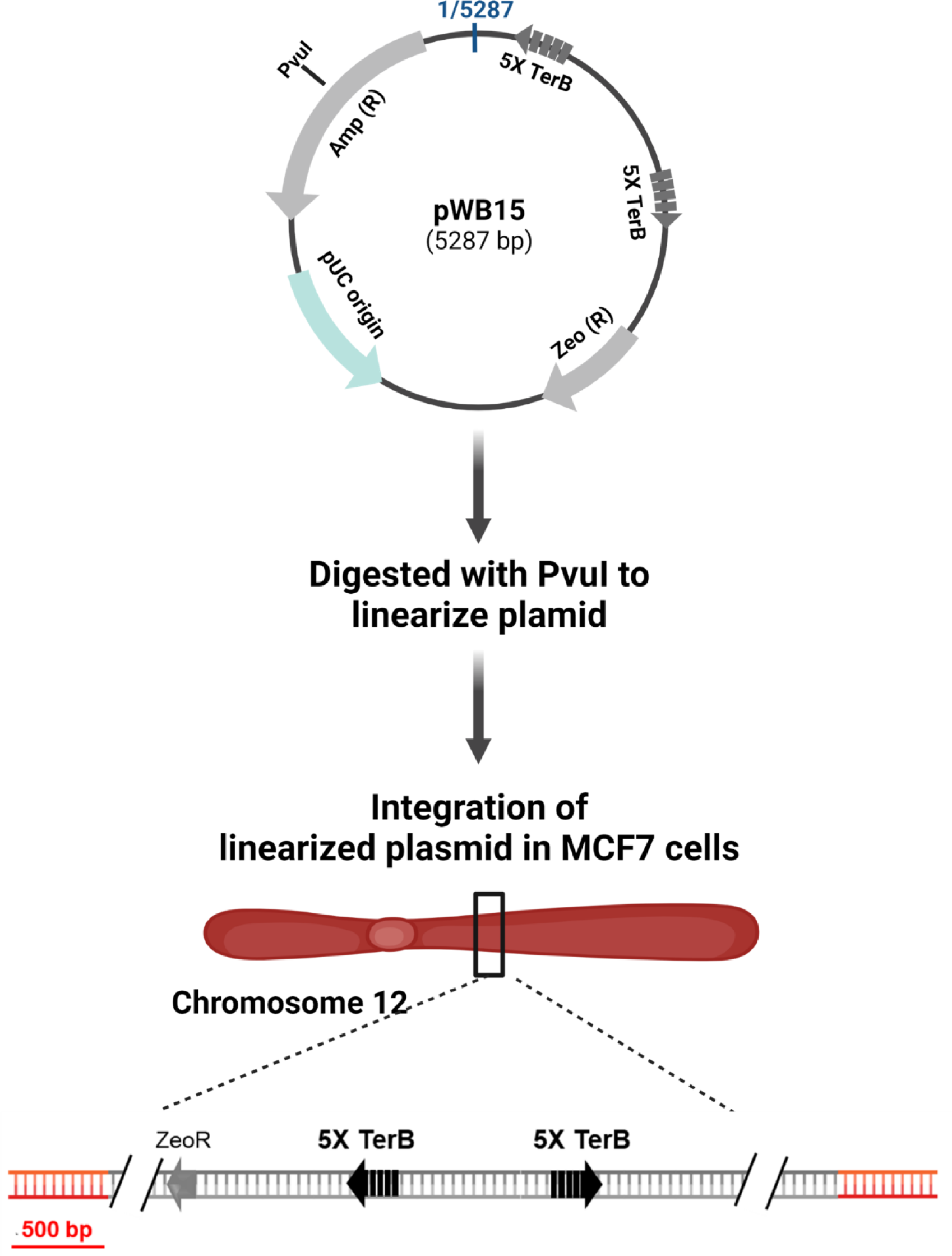
Generation of the MCF7 5C-TerB cell line. Schematic of the integration of pWB15 in MCF7 cells to generate MCF7 5C-TerB. pWB15 contains two *TerB* cassettes (grey arrows). Each *TerB* cassette contains 5 tandem *TerB* sequences, which are in the non-permissive orientation in pWB15 (grey arrow facing away from each other. The plasmid was linearized by digesting with PvuI for integration into MCF7 cells. The single copy integrant was confirmed using whole genome sequencing and found to be integrated at Chromosome 12.

**Supplementary Figure 2.**
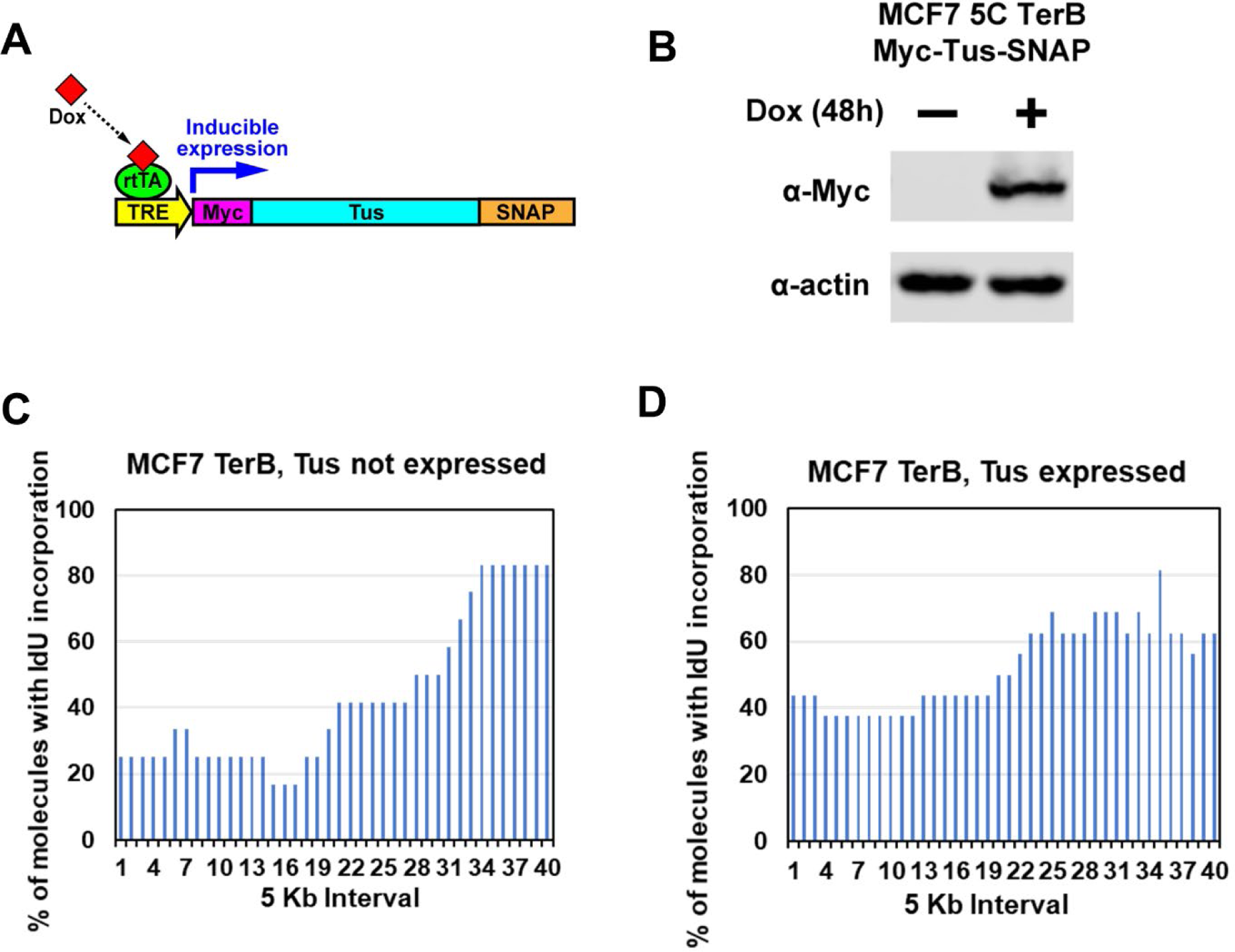
Expression of Tus in MCF7 5C-TerB cells. **(A)** Schematic of Doxycycline (Dox) inducible expression of Myc-NLS-TUS-SNAP. (rtTA = reverse tetracycline trans activator; TRE = tetracycline response element). **(B)** Immunoblot of MCF7 5C-TerB cells stably expressing Dox inducible Myc-NLS-TUS-SNAP. **(C-D)** Replication profiles shown as the percentage of molecules with IdU incorporation at each 5 Kb interval in the 200 Kb SfiI segment containing *TerB* sequence, quantified from molecules MCF7 5C-TerB in Figure 2C (Tus not expressed) and Figure 2D (Tus expressed).

**Supplementary Figure 3.**
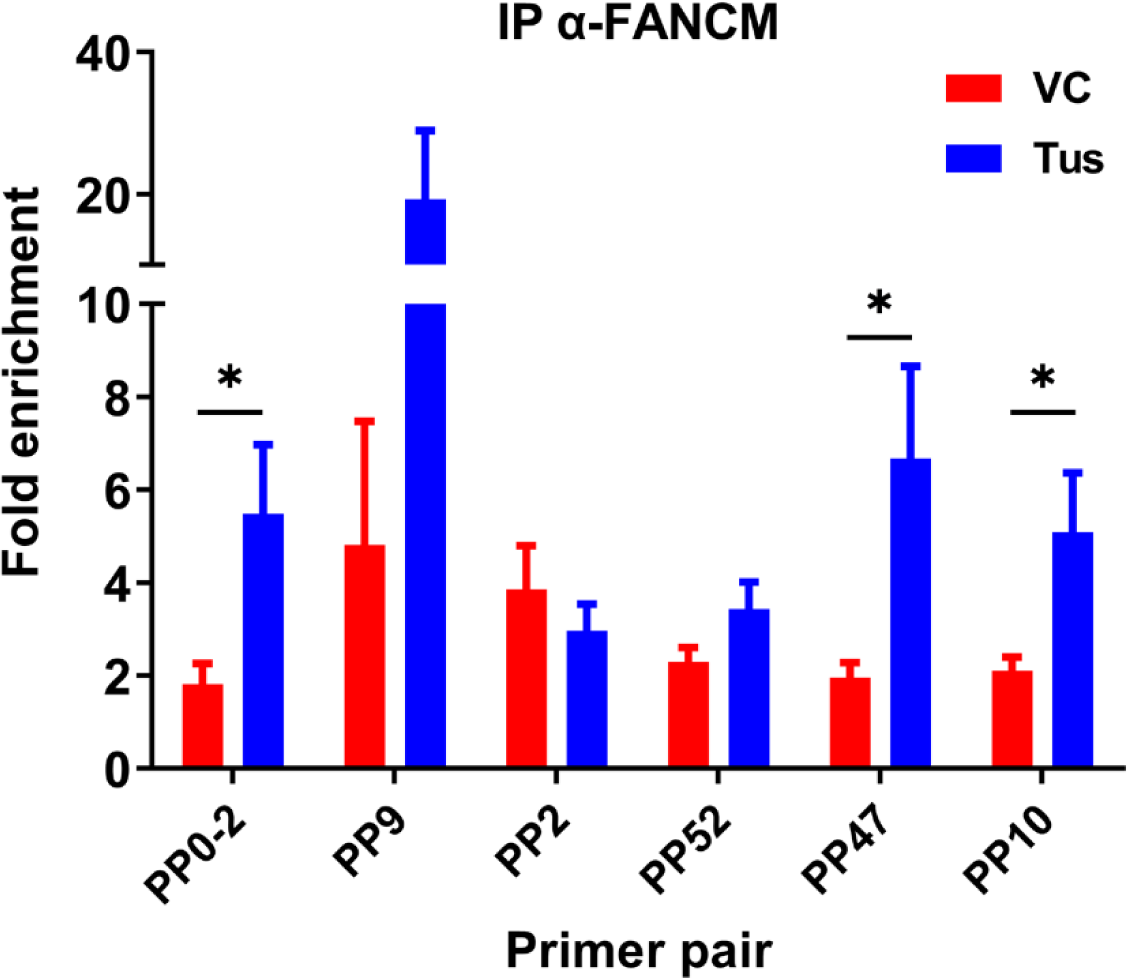
Enrichment of FANCM at the Ter sequence in the presence of the Tus protein. ChIP using FANCM antibody was performed MCF7 5C-TerB cells transfected with VC or HA-Tus-His plasmids. ChIP-qPCR were conducted using the indicated primer pairs. Data shows the fold enrichment relative to the IgG controls (n=3).

**Supplementary Figure 4.**
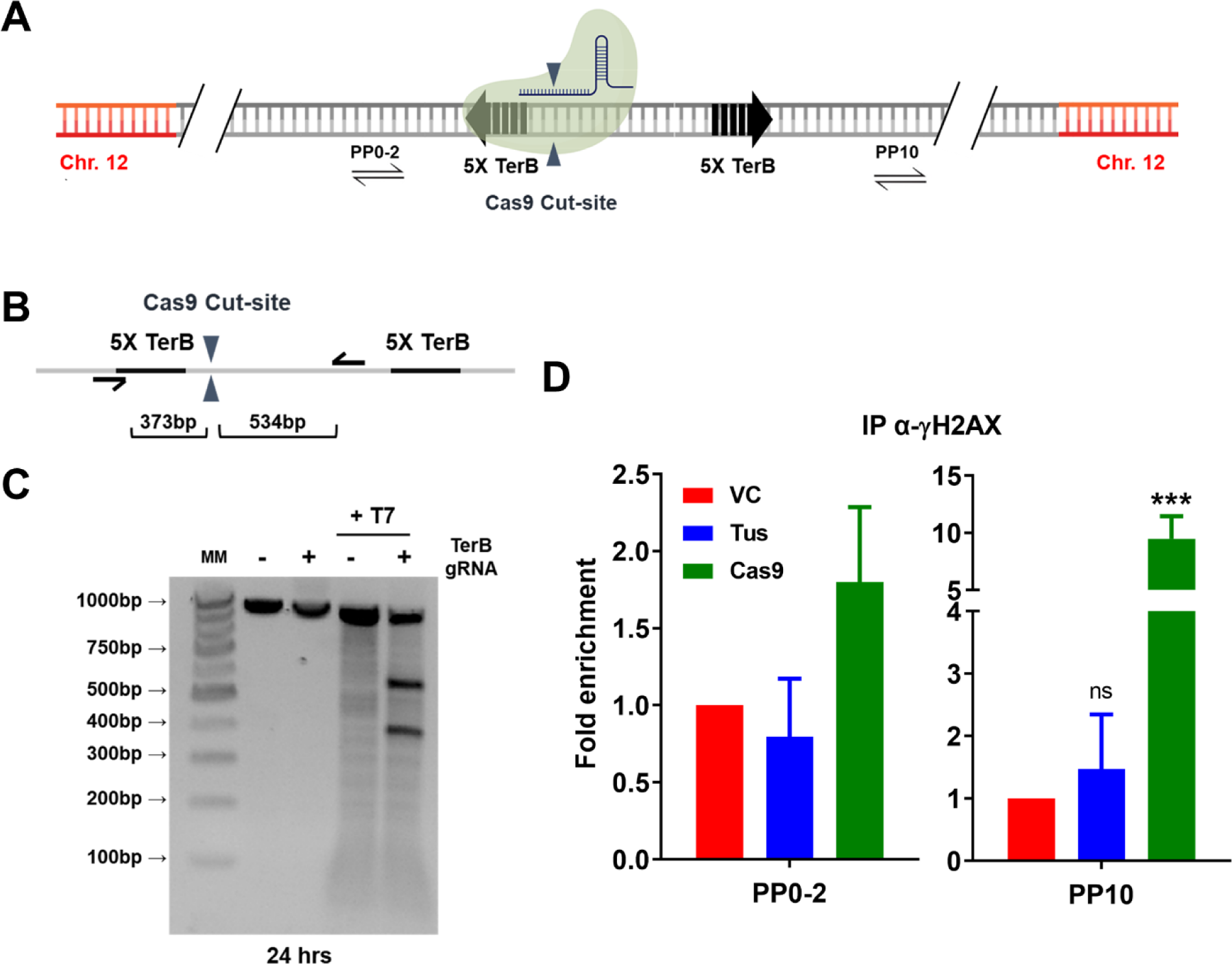
Distinct patterns of γH2AX enrichment at a Cas9-mediated DSB vs the Tus-*TerB* fork barrier. **(A)** Schematic of the linearized *TerB* plasmid (pWB15) integrated as a unique copy into MCF7 cells (MCF7 5C-TerB) with the Cas9-binding site (green protein). Blue triangles: Cas9 cut site. Black half-arrow heads depict PCR products expected from primer pair (PP10) used in quantitative PCR (qPCR) are shown. **(B)** Schematic depicting the Cas9 cut site (blue arrows), site-specific PCR primers (black half-arrowhead) and predicted size of cleavage products. **(C)** PCR analysis by T7 assay of MCF7 5C-TerB cells transfected with Cas9-sgRNA RNP complex. **(D)** γH2AX levels along the integrated *TerB* plasmid were analyzed by ChIP-qPCR in MCF7 5C-TerB cells transfected with VC, Tus or Cas9 expression plasmids using the indicated primer pairs. Data shows the fold enrichment relative to IgG controls (n=2).

**Supplementary Figure 5.**
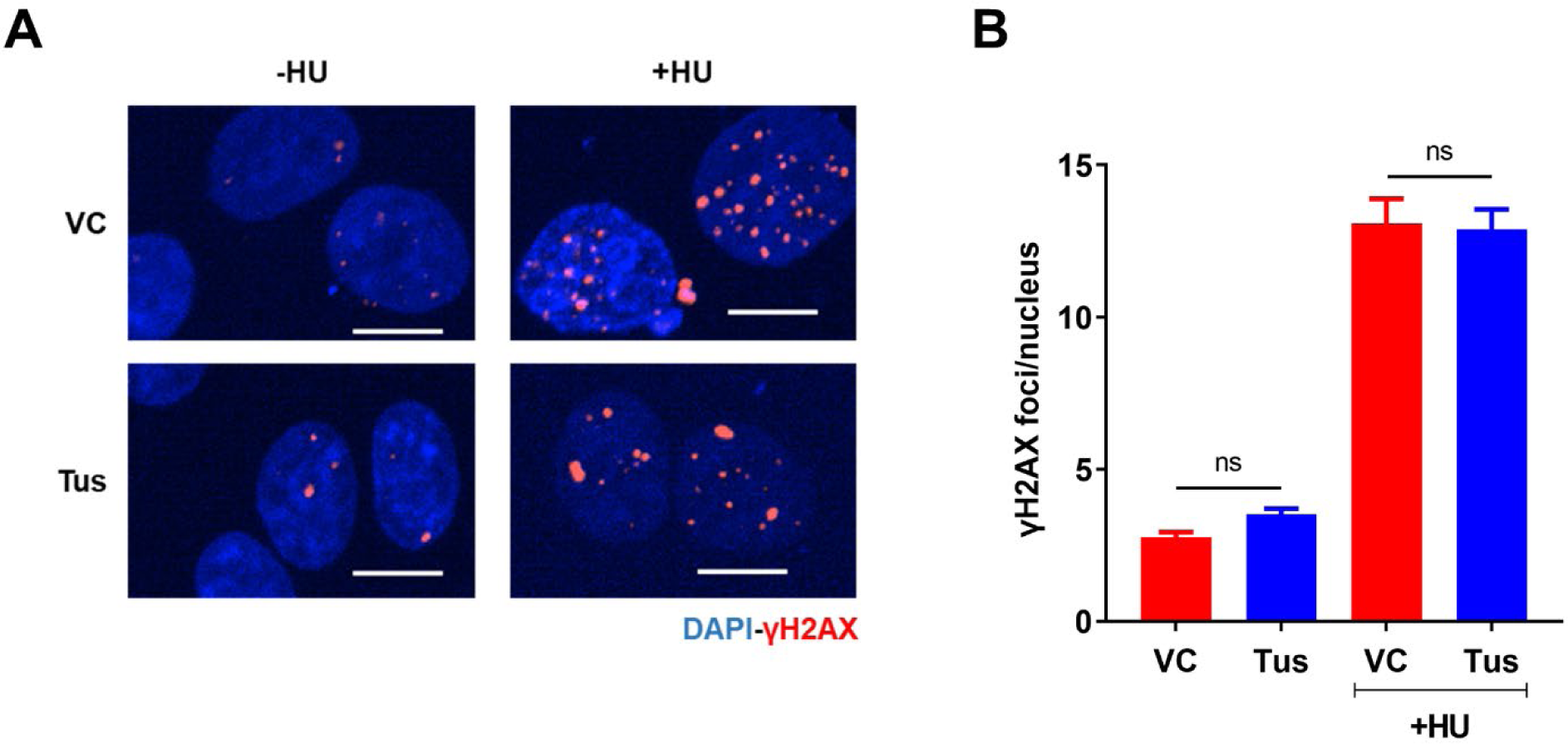
Genome-wide gH2AX foci were unaffected with Tus expression. **(A)** Representative images of gH2AX foci across stated conditions. Bars, 10 μm **(B)** Average number of gH2AX foci per nucleus in conditions stated. Cells treated with or without 2mM HU for 4 hours. (n=3, ≥300 cells per experiment)

**Supplementary Figure 6.**
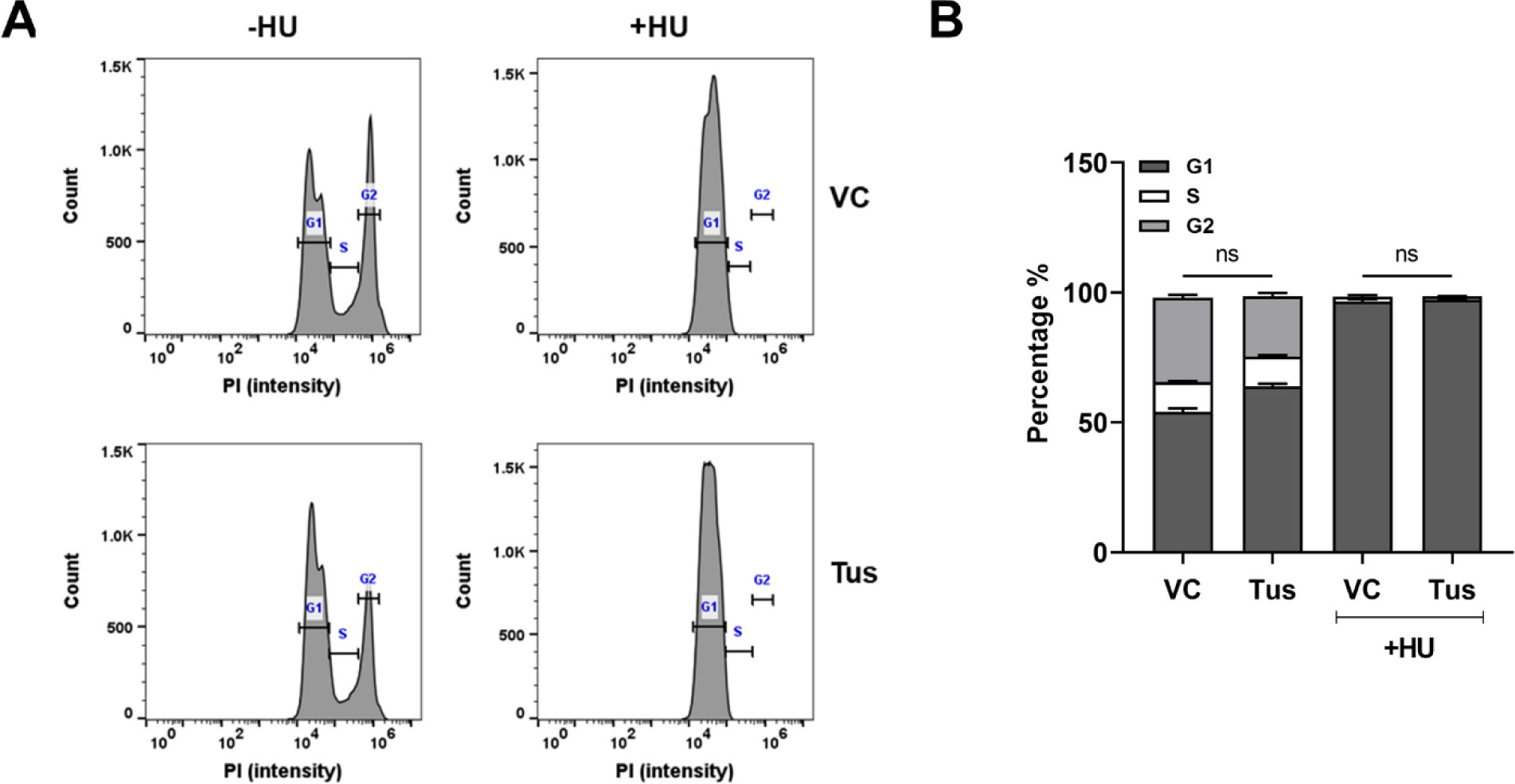
Cell cycle progression were unaffected with Tus expression. **(A)** Cell cycle analysis and **(B)** quantification was performed using flow cytometry following staining with propidium iodide staining in MCF7 5C-TerB cells under the stated conditions. Cells treated with or without 2mM HU for 4 hours. (n=3, ns: non-significant)

## Supplementary Tables

**Supplementary Table 1. Demonstration that the Tus-Ter replication fork block does not activate significant replication elsewhere in the genome.** Replication characteristics of 200 kb global DNA segments that represent the total genome. These measurements do not include the segments containing the *TerB* sequence. The table compares MCF7 cells containing the *TerB* sequence with and without Tus induced. This was determined on stretched DNA molecules that had completely incorporated IdU, CldU or a combination of both nucleotide analogues.

**Supplementary Table 2. List of antibodies, oligos and plasmids used in the study.**

## References

Bakkenist CJ, Kastan MB. 2003. DNA damage activates ATM through intermolecular autophosphorylation and dimer dissociation. Nature 2003 421:6922 421:499–506. doi:10.1038/nature01368

Ball HL, Myers JS, Cortez D. 2005. ATRIP binding to replication protein A-single-stranded DNA promotes ATR-ATRIP localization but is dispensable for Chk1 phosphorylation. Mol Biol Cell 16:2372–2381. doi:10.1091/MBC.E04-11-1006/ASSET/IMAGES/LARGE/ZMK0050531540006.JPEG

Bantele SCS, Lisby M, Pfander B. 2019. Quantitative sensing and signalling of single-stranded DNA during the DNA damage response. Nature Communications 2019 10:1 10:1–12. doi:10.1038/s41467-019-08889-5

Berkovich E, Monnat RJ, Kastan MB. 2007. Roles of ATM and NBS1 in chromatin structure modulation and DNA double-strand break repair. Nat Cell Biol 9:683–690. doi:10.1038/NCB1599

Brickner JR, Garzon JL, Cimprich KA. 2022. Walking a tightrope: The complex balancing act of R-loops in genome stability. Mol Cell 82. doi:10.1016/J.MOLCEL.2022.04.014

Bryan TM. 2019. Mechanisms of DNA Replication and Repair: Insights from the Study of G-Quadruplexes. Molecules 2019, *Vol* 24, *Page* 3439 24:3439. doi:10.3390/MOLECULES24193439

Chailleux C, Aymard F, Caron P, Daburon V, Courilleau C, Canitrot Y, Legube G, Trouche D. 2014. Quantifying DNA double-strand breaks induced by site-specific endonucleases in living cells by ligation-mediated purification. Nat Protoc 9:517–528. doi:10.1038/NPROT.2014.031

Chandramouly G, Kwok A, Huang B, Willis NA, Xie A, Scully R. 2013. BRCA1 and CtIP suppress long-tract gene conversion between sister chromatids. Nat Commun 4. doi:10.1038/NCOMMS3404

Clouaire T, Rocher V, Lashgari A, Arnould C, Aguirrebengoa M, Biernacka A, Skrzypczak M, Aymard F, Fongang B, Dojer N, Iacovoni JS, Rowicka M, Ginalski K, Côté J, Legube G. 2018. Comprehensive Mapping of Histone Modifications at DNA Double-Strand Breaks Deciphers Repair Pathway Chromatin Signatures. Mol Cell 72:250–262.e6. doi:10.1016/J.MOLCEL.2018.08.020

Collis SJ, Ciccia A, Deans AJ, Hořejší Z, Martin JS, Maslen SL, Skehel JM, Elledge SJ, West SC, Boulton SJ. 2008. FANCM and FAAP24 function in ATR-mediated checkpoint signaling independently of the Fanconi anemia core complex. Mol Cell 32:313–324. doi:10.1016/J.MOLCEL.2008.10.014

Dixon JR, Selvaraj S, Yue F, Kim A, Li Y, Shen Y, Hu M, Liu JS, Ren B. 2012. Topological domains in mammalian genomes identified by analysis of chromatin interactions. Nature 485:376–380. doi:10.1038/NATURE11082

Drosopoulos WC, Kosiyatrakul ST, Yan Z, Calderano SG, Schildkraut CL. 2012. Human telomeres replicate using chromosome-specific, rather than universal, replication programs. Journal of Cell Biology 197:253–266. doi:10.1083/JCB.201112083

Drosopoulos WC, Vierra DA, Kenworthy CA, Coleman RA, Schildkraut CL. 2020. Dynamic Assembly and Disassembly of the Human DNA Polymerase &delta; Holoenzyme on the Genome In&nbsp;Vivo. doi:10.1016/j.celrep.2019.12.101

Elango R, Panday A, Lach FP, Willis NA, Nicholson K, Duffey EE, Smogorzewska A, Scully R. 2022. The structure-specific endonuclease complex SLX4-XPF regulates Tus-Ter-induced homologous recombination. Nat Struct Mol Biol 29:801–812. doi:10.1038/S41594-022-00812-9

Fokas E, Prevo R, Pollard JR, Reaper PM, Charlton PA, Cornelissen B, Vallis KA, Hammond EM, Olcina MM, Gillies McKenna W, Musche RJ, Brunner TB. 2012. Targeting ATR in vivo using the novel inhibitor VE-822 results in selective sensitization of pancreatic tumors to radiation. Cell Death & Disease 2012 3:12 3:e441–e441. doi:10.1038/cddis.2012.181

Gaillard H, García-Muse T, Aguilera A. 2015. Replication stress and cancer. Nat Rev Cancer 15:276–280. doi:10.1038/NRC3916

Helt CE, Wang W, Keng PC, Bambara RA. 2005. 9-1-1 Complex Involvement in DNA Repair: Evidence that DNA Damage Detection Machinery Participates in DNA Repair. http://dx.doi.org/104161/cc4415984:529-532. doi:10.4161/CC.4.4.1598

Hiasa H, Marians KJ. 1994. Tus prevents overreplication of oriC plasmid DNA. Journal of Biological Chemistry 269:26959–26968. doi:10.1016/S0021-9258(18)47112-8

Hidaka M, Akiyama M, Horiuchi T. 1988. A consensus sequence of three DNA replication terminus sites on the E. coli chromosome is highly homologous to the terR sites of the R6K plasmid. Cell 55:467–475. doi:10.1016/0092-8674(88)90033-5

Iyer DR, Rhind N. 2017. The Intra-S Checkpoint Responses to DNA Damage. Genes (Basel) 8. doi:10.3390/GENES8020074

Iyer DR, Rhind N. 2013. Checkpoint regulation of replication forks: global or local? Biochem Soc Trans 41:1701–1705. doi:10.1042/BST20130197

Janke R, Herzberg K, Rolfsmeier M, Mar J, Bashkirov VI, Haghnazari E, Cantin G, Yates JR, Heyer WD. 2010. A truncated DNA-damage-signaling response is activated after DSB formation in the G1 phase of Saccharomyces cerevisiae. Nucleic Acids Res 38:2302. doi:10.1093/NAR/GKP1222

Kaufmann WK, Cleaver JE, Painter RB. 1980. Ultraviolet radiation inhibits replicon initiation in S phase human cells. Biochim Biophys Acta 608:191–195. doi:10.1016/0005-2787(80)90147-1

Kim JC, Harris ST, Dinter T, Shah KA, Mirkin SM. 2016. The role of break-induced replication in large-scale expansions of (CAG)n/(CTG)n repeats. Nature Structural & Molecular Biology 2016 24:1 24:55–60. doi:10.1038/nsmb.3334

Larsen NB, Hickson ID, Mankouri HW. 2014a. Tus-Ter as a tool to study site-specific DNA replication perturbation in eukaryotes. Cell Cycle 13:2994. doi:10.4161/15384101.2014.958912

Larsen NB, Sass E, Suski C, Mankouri HW, Hickson ID. 2014b. The Escherichia coli Tus– Ter replication fork barrier causes site-specific DNA replication perturbation in yeast. Nature Communications 2014 5:1 5:1–10. doi:10.1038/ncomms4574

Marie L, Symington LS. 2022. Mechanism for inverted-repeat recombination induced by a replication fork barrier. Nature Communications 2022 13:1 13:1–13. doi:10.1038/s41467-021-27443-w

Merrick CJ, Jackson D, Diffley JFX. 2004. Visualization of altered replication dynamics after DNA damage in human cells. J Biol Chem 279:20067–20075. doi:10.1074/JBC.M400022200

Mirkin E v., Mirkin SM. 2007. Replication Fork Stalling at Natural Impediments. Microbiol Mol Biol Rev 71:13. doi:10.1128/MMBR.00030-06

Mulcair MD, Schaeffer PM, Oakley AJ, Cross HF, Neylon C, Hill TM, Dixon NE. 2006. A molecular mousetrap determines polarity of termination of DNA replication in E. coli. Cell 125:1309–1319. doi:10.1016/J.CELL.2006.04.040

Nandi S, Whitby MC. 2012. The ATPase activity of Fml1 is essential for its roles in homologous recombination and DNA repair. Nucleic Acids Res 40:9584–9595. doi:10.1093/NAR/GKS715

Norio P, Schildkraut CL. 2001. Visualization of DNA replication on individual Epstein-Barr virus episomes. Science 294:2361–2364. doi:10.1126/SCIENCE.1064603

Panday A, Willis NA, Elango R, Menghi F, Duffey EE, Liu ET, Scully R. 2021. FANCM regulates repair pathway choice at stalled replication forks. Mol Cell 81:2428–2444.e6. doi:10.1016/J.MOLCEL.2021.03.044/ATTACHMENT/DB09DDDE-5057-4829-A9A6-753D68F31F16/MMC1.PDF

Parrilla-Castellar ER, Arlander SJH, Karnitz L. 2004. Dial 9–1–1 for DNA damage: the Rad9–Hus1–Rad1 (9–1–1) clamp complex. DNA Repair (Amst) 3:1009–1014. doi:10.1016/J.DNAREP.2004.03.032

Rao SSP, Huntley MH, Durand NC, Stamenova EK, Bochkov ID, Robinson JT, Sanborn AL, Machol I, Omer AD, Lander ES, Aiden EL. 2014. A 3D map of the human genome at kilobase resolution reveals principles of chromatin looping. Cell 159:1665–1680. doi:10.1016/J.CELL.2014.11.021

Roecklein B, Pelletier A, Kuempel P. 1991. The tus gene of Escherichia coli: autoregulation, analysis of flanking sequences and identification of a complementary system in Salmonella typhimurium. Res Microbiol 142:169–175. doi:10.1016/0923-2508(91)90026-7

Savic V, Yin B, Maas NL, Bredemeyer AL, Carpenter AC, Helmink BA, Yang-Iott KS, Sleckman BP, Bassing CH. 2009. Formation of dynamic gamma-H2AX domains along broken DNA strands is distinctly regulated by ATM and MDC1 and dependent upon H2AX densities in chromatin. Mol Cell 34:298–310. doi:10.1016/J.MOLCEL.2009.04.012

Saxena S, Zou L. 2022. Hallmarks of DNA replication stress. Mol Cell 82:2298–2314. doi:10.1016/J.MOLCEL.2022.05.004

Shimada K, Pasero P, Gasser SM. 2002. ORC and the intra-S-phase checkpoint: a threshold regulates Rad53p activation in S phase. Genes Dev 16:3236–3252. doi:10.1101/GAD.239802

Tubbs A, Nussenzweig A. 2017. Endogenous DNA Damage as a Source of Genomic Instability in Cancer. Cell 168:644–656. doi:10.1016/J.CELL.2017.01.002

Twayana S, Bacolla A, Barreto-Galvez A, De-Paula RB, Drosopoulos WC, Kosiyatrakul ST, Bouhassira EE, Tainer JA, Madireddy A, Schildkraut CL. 2021. Translesion polymerase eta both facilitates DNA replication and promotes increased human genetic variation at common fragile sites. Proc Natl Acad Sci U S A 118:e2106477118. doi:10.1073/PNAS.2106477118/SUPPL_FILE/PNAS.2106477118.SD03.XLSX

Ward IM, Chen J. 2001. Histone H2AX Is Phosphorylated in an ATR-dependent Manner in Response to Replicational Stress. Journal of Biological Chemistry 276:47759– 47762. doi:10.1074/jbc.C100569200

Willis N, Rhind N. 2009a. Regulation of DNA replication by the S-phase DNA damage checkpoint. Cell Div 4:1–10. doi:10.1186/1747-1028-4-13/COMMENTS

Willis N, Rhind N. 2009b. Regulation of DNA replication by the S-phase DNA damage checkpoint. Cell Div 4. doi:10.1186/1747-1028-4-13

Willis NA, Chandramouly G, Huang B, Kwok A, Follonier C, Deng C, Scully R. 2014. BRCA1 controls homologous recombination at Tus/Ter-stalled mammalian replication forks. Nature 510:556–559. doi:10.1038/NATURE13295

Willis NA, Frock RL, Menghi F, Duffey EE, Panday A, Camacho V, Hasty EP, Liu ET, Alt FW, Scully R. 2017. Mechanism of tandem duplication formation in BRCA1-mutant cells. Nature 2017 551:7682 551:590–595. doi:10.1038/nature24477

Willis NA, Panday A, Duffey EE, Scully R. 2018. Rad51 recruitment and exclusion of non-homologous end joining during homologous recombination at a Tus/Ter mammalian replication fork barrier. PLoS Genet 14. doi:10.1371/JOURNAL.PGEN.1007486

Willis NA, Scully R. 2016. Spatial separation of replisome arrest sites influences homologous recombination quality at a Tus/Ter-mediated replication fork barrier. Cell Cycle 15:1812–1820. doi:10.1080/15384101.2016.1172149

Xue X, Sung P, Zhao X. 2015. Functions and regulation of the multitasking FANCM family of DNA motor proteins. Genes Dev 29:1777–1788. doi:10.1101/GAD.266593.115

Zeman MK, Cimprich KA. 2013. Causes and consequences of replication stress. Nature Cell Biology 2014 16:1 16:2–9. doi:10.1038/ncb2897

Zou L, Elledge SJ. 2003. Sensing DNA damage through ATRIP recognition of RPA-ssDNA complexes. Science (1979) 300:1542–1548. doi:10.1126/SCIENCE.1083430/SUPPL_FILE/ZOU.SOM.PDF

