## Supplemental Tables for "A local ATR-dependent checkpoint pathway is activated by a site-specific replication fork block in human cells"

### Supplementary Table 1

| | **MCF7 with integrated *TerB*** | **MCF7 with integrated *TerB* + Tus expressed** |
|---|---|---|
| Total number of 200 kb DNA segments counted | 291 | 334 |
| Number of segments containing both red and green signal (RG) | 85 | 108 |
| Time to replicate (minutes) | 87 | 92 |
| Fork Rate (kb/min) per (RG) | 1.7 | 1.6 |

**Supplementary Table 1.Demonstration that the Tus-Ter replication fork block does not activate significant replication elsewhere in the genome**. Replication characteristics of 200 kb global DNA segments that represent the total genome. These measurements do not include the segments containing the *TerB* sequence. The table compares MCF7 cells containing the *TerB* sequence with and without Tus induced. This was determined on stretched DNA molecules that had completely incorporated IdU, CldU or a combination of both nucleotide analogues**.**

### Supplementary Table 2

| | Manufacturer | Catalogue No. |
|---|---|---|
| **Antibodies** | | |
| Actin | Sigma | A2066 |
| Avidin, NeutrAvidin™, Alexa Fluor™ 350 conjugate | Invitrogen | A11236 |
| Anti-Mouse IgG, HRP-linked | Cell signaling | #7076 |
| Anti-Rabbit IgG, HRP-linked | Cell signaling | #7074 |
| ATR Th1989 | Genetex | GTX128145 |
| Chk1 S345 | Cell signaling | #2348 |
| FANCM | Abcam | Ab95014 |
| GAPDH FL-335 | Santa Cruz | sc-25778 |
| GFP | Abcam | Ab290 |
| GFP B-2 | Santa Cruz | SC 9996 |
| gH2AX s139 | Abcam | Ab2893 |
| gH2AX s139 | Abcam | Ab81299 |
| gH2AX s139 | Cell Signaling | 9718S |
| Goat anti-Rabbit, Alexa Fluor™ Plus 555 | ThermoFisher Scientific | A32732 |
| Goat anti-Rat IgG, Alexa Fluor™ 488 | Invitrogen | A-11006 |
| Goat anti-Mouse IgG, Alexa Fluor™ 568 | Invitrogen | A-11031 |
| Goat Anti-Mouse IRDye 680LT | LiCor | #926-68020 |
| Goat Anti-Rabbit IRDye 800CW | LiCor | #926-32211 |
| Goat Anti-Avidin D Antibody, Biotinylated | Vector Laboratories | BA-0300 |
| HA F-7 | Santa Cruz | sc-7392 |
| His | Abcam | Ab9108 |
| IgG | Cell signaling | #2729 |
| Lamin A/C | Santa Cruz | sc-6215 |
| MCM3 | Abcam | Ab4460 |
| Myc | Cell Signaling | #2276 |

| | | |
|---|---|---|
| Purified Mouse Anti-BrdU | BD Biosciences | 347580 |
| Rat monoclonal anti-BrdU antibody | Abcam | ab6326 |
| RPA 32 S33 | Bethyl | A300-246 |
| Total ATR (N-19) | Santa Cruz | sc-1887 |
| Total Chk1 (G-4) | Santa Cruz | sc-8408 |
| Total RPA 32 | Cell Signaling | #2208 |
| Tus | This study | NA |
| **Oligonucleotides** | | |
| Primers for PCR amplification of pcDNA3-β-MYC-NLS-Tus,<br><br>Forward: AGTCGGTACCGAATTCGCCACCATGGAACAAAAGCTG<br><br>Reverse: AGTCGGCGGCCGCGCCGCTACCGTCAGCCACGTACAGGTGCA | This paper | N/A |
| Primers for PCR amplification of SNAP tag cDNA,<br><br>Forward: AGTCGCGGCCGCCGGCCACATGGACAAAGACTGCGAAATGAAGC<br><br>Reverse: ACTGCTCGAGTCAACCCAGCCCAGGCTTGC | This paper | N/A |
| sgRNA TerB1:TTGCGCTGCTTCGCGATGTA | This paper | N/A |
| Primer Pair PP-0-2 Forward: TCTGAGAATAGTGTATGCGG Reverse: AGATGCTGAAGATCAGTTGG | This paper | N/A |
| Primer Pair PP9<br><br>Forward: CGAGCTCGGATCAATAAGT<br><br>Reverse: AGAGTCGACCATAGGGGAT | This paper | N/A |
| Primer Pair PP2<br><br>Forward: AAAGTTCGAGTCTAGAGGGC<br><br>Reverse: GCATCAGAGCAGATTGTACT | This paper | N/A |

| | | |
|---|---|---|
| Primer Pair PP52<br>Forward: TCCTACTTGGCAGTACATCT<br>Reverse: GGAAAGTCCCGTTGATTTTG | This paper | N/A |
| Primer Pair PP47<br>Forward: AGCGTTTAAACTTAAGCTTGGTA<br>Reverse: GGCCCTCTAGACTCGAAATAA | This paper | N/A |
| Primer Pair PP10<br>Forward: TTTAGGGTTCCGATTTAGTGCT<br>Reverse: ATTTTTTAACCAATAGGCCGA | This paper | N/A |
| **Plasmids** | | |
| Myc-NLS-TUS-SNAP | This study | N/A |
| pcDNA3-β-MYC-NLS-Tus | This study | N/A |
| pCMV3xnls | Lab stock | N/A |
| pCMV3xnls-HA | This study | N/A |
| pCMV3xnls-GFP | This study | N/A |
| pCMV3xnls-Tus | This study | N/A |
| pCMV3xnls-Tus-His | This study | N/A |
| pCMV3xnls-Tus-HA | This study | N/A |
| pCMV3xnls-Tus-GFP | This study | N/A |
| pWB15 | This study | N/A |
| pInd Tus-SNAP | This study | N/A |
| pInducer10L | Drosopoulos et al., Cell Reports 30 2020 | N/A |
| Fosmid | BACPAC Genomic | WI2-1478M20 |
